# Synergistic targeting of MCL1 and caspases for enhanced anti-tumor immunity in breast cancer

**DOI:** 10.1101/2025.10.10.681458

**Authors:** Jae Kyo Yi, Ji-Sun Lee, Pilhan Kim, Leslie M. Shaw, Alexander Spektor

## Abstract

Breast cancer remains largely unresponsive to immunotherapy due to its immunologically “cold” nature, marked by low cytolytic activity and a paucity of neoantigens. To overcome this, we developed a novel therapeutic approach to activate the cGAS-STING pathway, a critical mediator of the type I interferon response essential for effective anti-tumor immunity. We demonstrate that combined inhibition of MCL1 and caspases robustly triggers a type I IFN response in breast cancer cells, effectively remodeling the tumor microenvironment by increasing immune cell infiltration and enhancing antigen presentation. In immunocompetent syngeneic mouse models, this combination therapy significantly suppressed tumor growth, an effect that was reversed upon blockade of the IFN signaling axis. Mechanistically, our findings reveal that co-targeting MCL1 and caspases reprograms the tumor microenvironment enhancing immune surveillance. Given the established safety profiles of both drug classes, this strategy offers a promising and rapidly deployable approach to sensitize breast tumors to immunotherapy.

---

Breast cancer, especially the estrogen receptor-positive (ER+) subtype, has long been characterized by its immunologically “cold” nature, a term that describes tumors with low immune cytolytic activity and a paucity of neo-epitopes due to a relatively small number of nonsynonymous mutations^1, 2^. This immunological profile poses significant challenges for immune-based therapeutic strategies, as the lack of neo-epitopes results in diminished recognition and eradication by the host immune system ^3, 4^. Consequently, there is a need for novel strategies that can enhance the immunogenicity of breast cancers.

Recent studies have extensively focused on utilizing the cGAS-STING pathway, a critical mediator of the type I interferon (IFN) response pivotal in anti-tumor immunity, to favorably alter the tumor microenvironment for cancer therapy ^5, 6^. The activation of this pathway typically results in upregulation of type I IFNs, which can bolster immune surveillance and potentially counteract tumor immune evasion ^7^. However, emerging evidence suggests that in the context of chromosomally unstable cancers, there is a paradoxical effect. Chronic activation of cGAS, accompanied by STING downregulation—a phenomenon known as tachyphylaxis—and induction of endoplasmic reticulum (ER) stress, leads not to immunostimulation but rather to a state of immunosuppression ^8^. This immunosuppressive milieu is further exacerbated by the promotion of a pro-inflammatory response that fosters metastasis ^9^. In this landscape, conventional treatments aiming to amplify the IFN response, such as STING agonists, fail to achieve their intended effects, thus necessitating the exploration of alternative approaches ^10, 11^. One recently proposed strategy is to target damage-associated molecular patterns (DAMPs), such as oxidized proteins and mitochondrial DNA (mtDNA), which are capable of inducing an IFN response ^12–14^. However, the effectiveness of DAMPs is contingent upon their release from the mitochondria, where they are typically sequestered ^15–17^. Apoptotic signaling leading to mitochondrial outer membrane permeabilization (MOMP) facilitates the release of these DAMPs, enabling them to engage pattern recognition receptors (PRRs) that initiate the IFN response ^18^. In physiological conditions, the regulation of MOMP is tightly controlled by members of the BCL2 family of proteins ^19^. However, MCL1, a key BCL2 family member, is often dysregulated in breast cancer ^20, 21^. Interestingly, breast cancer cells exhibit significant dependence on MCL1, making it a promising target for therapeutic intervention ^22^. Indeed, inhibitors of MCL1 have shown efficacy in early clinical trials, indicating their potential role in inducing MOMP and thereby promoting a type I IFN response specifically within cancer cells ^23,24^.

In this study, we demonstrated that strategically inducing MOMP through MCL1 inhibition, combined with the suppression of caspases—key enzymes known to cleave and inactivate PRRs—can elicit a robust and targeted type I IFN response ^25, 26^. This dual inhibition approach effectively reshaped the tumor microenvironment, creating conditions conducive to a heightened anti-tumor immune response. Our findings reveal the significant impact of enhanced type I IFN signaling on both the tumor microenvironment and immune-mediated tumor control, supported by *in vitro* and *in vivo* evidence. These results present a promising therapeutic strategy to remodel the tumor microenvironment and bolster anti-tumor immunity with strong potential for rapid clinical application.

## Results

### Type I IFN response is induced by the dual inhibition of MCL1 and caspases in ER+ breast cancer cell lines

Studies have endeavored to potentiate the type I IFN response as a strategy to modulate the tumor microenvironment and augment tumor control ^11, 27^. Various anti-cancer drugs targeting proteins like BCL2, CDK4/6, and PGC1a have demonstrated effectiveness in their primary functions and additionally shown potential in inducing IFN responses in cancer cells ^28–30^. Our study thus examined the effects of drugs including ribociclib (CDK4/6 inhibitor), AZD4320 (BCL2 inhibitor), SR18292 (PGC1a modulator), and AZD5991 (MCL1 inhibitor) on the activation of type I IFN genes IFNB1, OAS1 and IFIT1 in MCF-7 cells. The integrity of the IFN response in MCF-7 cells was confirmed by treatment with nucleic acid analogs dA:dT and Poly I:C, which induced significant expression of both genes (Fig. 1a). Among the drugs tested, only AZD5991 (MCL1 inhibitor) induced a moderate increase in the expression levels of IFNB1 and OAS1 (Fig. 1a). Subsequently, we explored whether MCL1 inhibition could trigger a type I IFN response in different breast cancer cell lines. Although MCL1 inhibitors demonstrated comparable cytotoxic effects in MCF-7, ZR-75-1, and T47d cells (Extended Data Fig. 1a), ZR-75-1 and T47d did not elicit the same IFN response as MCF-7 (Fig. 1b).

**Figure 1:**
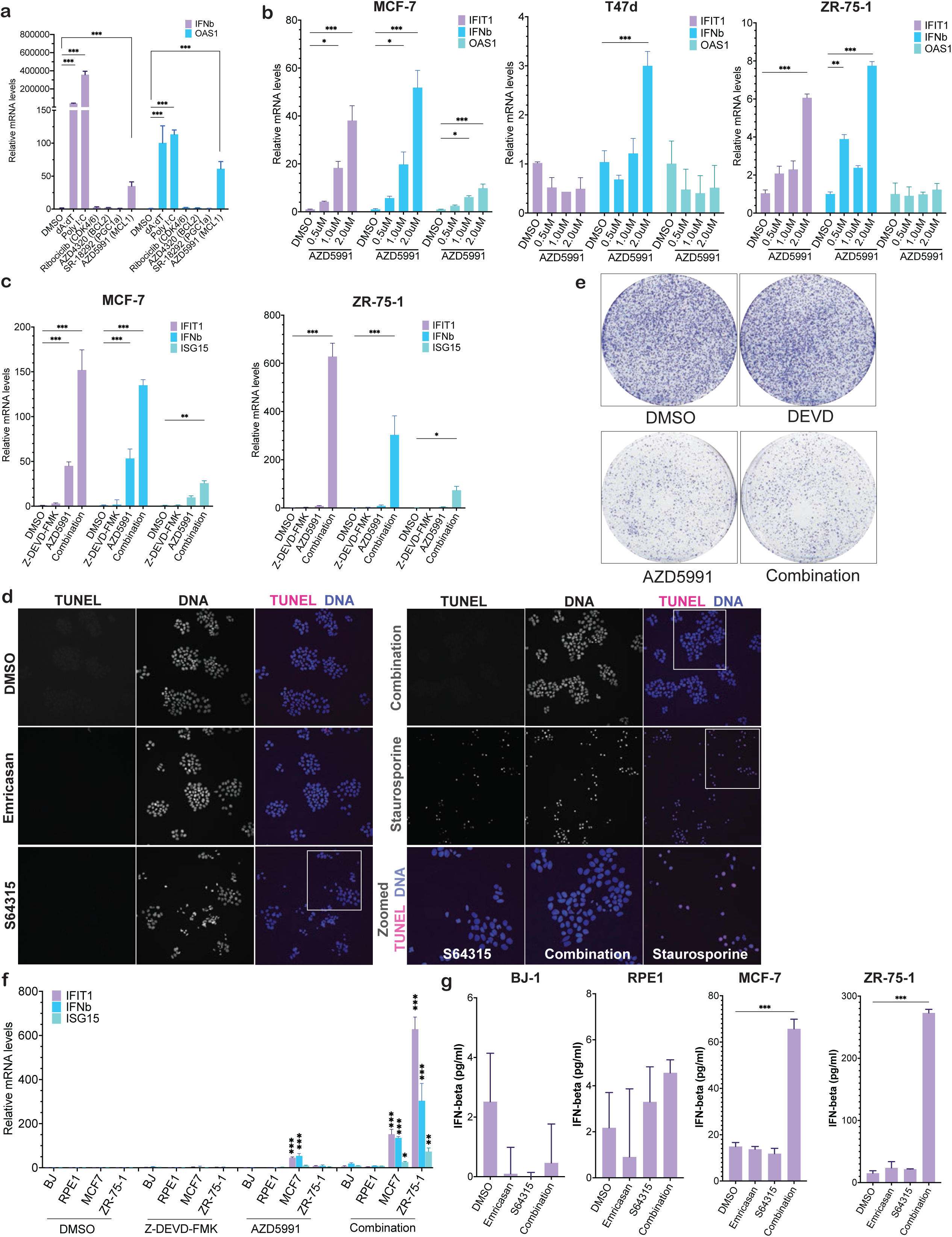
Differential induction of type I IFN response by MCL1 and caspase dual inhibition in ER+ breast cancer cell lines. **a,** Quantitative PCR analysis of IFNB1 and OAS1 gene expression in MCF-7 cells. Synthetic nucleic acid analogs dA:dT and Poly I:C, served as positive controls for IFN response activation. Cells were treated with anti-cancer agents targeting BCL2 (1 µM AZD4320), CDK4/6 (1µM Ribociclib), and PGC1a (5 µM SR18292). Data represent mean +/- SEM; unpaired t-test; n=3 biologically independent samples. **b,** Quantitative PCR of type I IFN genes across breast cancer cell lines (MCF-7, ZR-75-1 and T47d) upon MCL1 inhibition with 2 µM AZD5991. Data represent mean +/- SEM; unpaired t-test; n=3 biologically independent samples. **c,** Quantitative PCR to measure expression of IFNB1 and ISGs (IFIT1 and ISG15) upon combined treatment with MCL1 inhibitor AZD5991 (2 µM) and caspase-3 inhibitor Z-DEVD-FMK (10 µM) in ER+ breast cancer cell lines (MCF-7 and ZR-75-1). **d**, TUNEL assay to detect apoptosis of ZR-75-1 upon combined treatment with MCL1 inhibitor S64315 (2 µM) and caspase-3 inhibitor emricasan (10 µM) or staurosporine (positive control, 1 µM) **e,** Colony formation assay of ZR-75-1 cells under treatment with MCL1 inhibitor AZD5991 alone (2 µM) or in combination with caspase inhibitor (10 µM). (n=3 biologically independent samples) **f and g,** Analysis of type I IFN response by qPCR **(f)** and ELISA **(g)** for IFNB1 expression levels or protein levels in the media, respectively. ER+ breast cancer cell lines and non-transformed cell lines (BJ-hTERT and RPE1-hTERT) were treated with MCL1 inhibitor S64315 (2 µM) and/or caspase-3 inhibitor emricasan (10 µM). Data represent mean ± SEM; unpaired t-test; n=3 biologically independent samples. Statistical significance is indicated as *p < 0.05, **p < 0.01, ***p < 0.001.

Given that type I IFN response is inhibited by caspase-mediated cleavage of PRRs ^26^ and MCF-7 is deficient for caspase-3 ^31^, we hypothesized that combining MCL1 inhibitors with caspase inhibitors would greatly enhance type I IFN response in a robust and specific manner by facilitating release on DAMPs into the cytoplasm while preventing inactivation of PRRs. Strikingly, in contrast with MCL1 inhibitor treatment alone, combination treatment with Z-DEVD-FMK (caspase-3 inhibitor) led to a dramatic increase in expression levels of IFNB1 and the interferon-stimulated genes (ISGs), IFIT1 and ISG15 in both MCF-7 and ZR-75-1 ER+ breast cancer cell lines (Fig. 1c). However, this combination treatment did not significantly alter the cytotoxic effects compared to AZD5991 treatment alone (Extended Data Fig. 1b).

To verify these findings, we first assessed apoptosis using a TUNEL assay. Intriguingly, apoptosis was not observed after treatment with Emricasan (pan-caspase inhibitor) ^32^ or S64315 (MCL1 inhibitor) ^33^, and was only induced by the positive control, staurosporine (Fig. 1d). Furthermore, we conducted a colony forming assay on ZR-75-1 cells. While MCL1 inhibitor markedly decreased colony numbers, there was no discernible difference from the combination treatment (Fig. 1e). These results suggest that the MCL1 and caspase inhibitor combination can induce an IFN response at concentrations that do not cause apoptosis. Additionally, the addition of a caspase inhibitor does not affect cytotoxicity or alter clonogenic capacity following MCL1 inhibition, but it does induce a robust type I IFN response when combined with MCL1 inhibition. Interestingly, treatment with MCL1 inhibitor did not cause cytotoxicity, and the combination treatment minimally induced an IFN response in another breast cancer cell line, T47d (Extended Data Fig. 1c). Given that T47d has been reported to be resistant to MCL1 inhibitor treatment due to the high baseline level of another prosurvival protein, BCL2 ^34, 35^, we investigated whether simultaneous inhibition of BCL2 and MCL1 could trigger an IFN response in T47d when combined with caspase inhibitor. Indeed, triple inhibition of BCL2, MCL1 and caspases markedly increased the type I IFN response and cytotoxicity in T47d cells (Extended Data Fig. 1d and 1e), suggesting that both effective MOMP and simultaneous caspase inhibition are needed to elicit a robust type I IFN response.

To assess the effects of dual caspase and MCL1 inhibition on non-transformed cells, we also evaluated the impact of the combination on IFN response in BJ-hTERT (fibroblast) and RPE1-hTERT (epithelial cells) cell lines. Strikingly, the combination treatment induced only a minimal type I IFN response, as demonstrated by both mRNA expression levels and IFN-β concentrations (Fig. 1f and 1g), suggesting that IFN response to this drug combination is limited to cancer cells.

### The combination treatment positively alters the immunological landscape *in vitro*

To investigate alterations in the transcriptomic landscape from the combination treatment, RNA-sequencing (RNA-seq) was performed on ZR-75-1 cells treated with MCL1 inhibitor and/or caspase-3 inhibitor. Gene Set Enrichment Analysis (GSEA) was conducted against the “Hallmark” gene sets to understand the impact on cellular pathways (Fig. 2a, Extended Data Fig 2a). Cells treated with the combination treatment demonstrated marked enrichment of innate immune response pathways, cytokine and chemokine activity and antigen processing and presentation, supporting our hypothesis that dual targeting of MCL1 and caspases regulates the immune landscape of breast cancer cells, potentially enhancing their immunogenicity.

**Figure 2.**
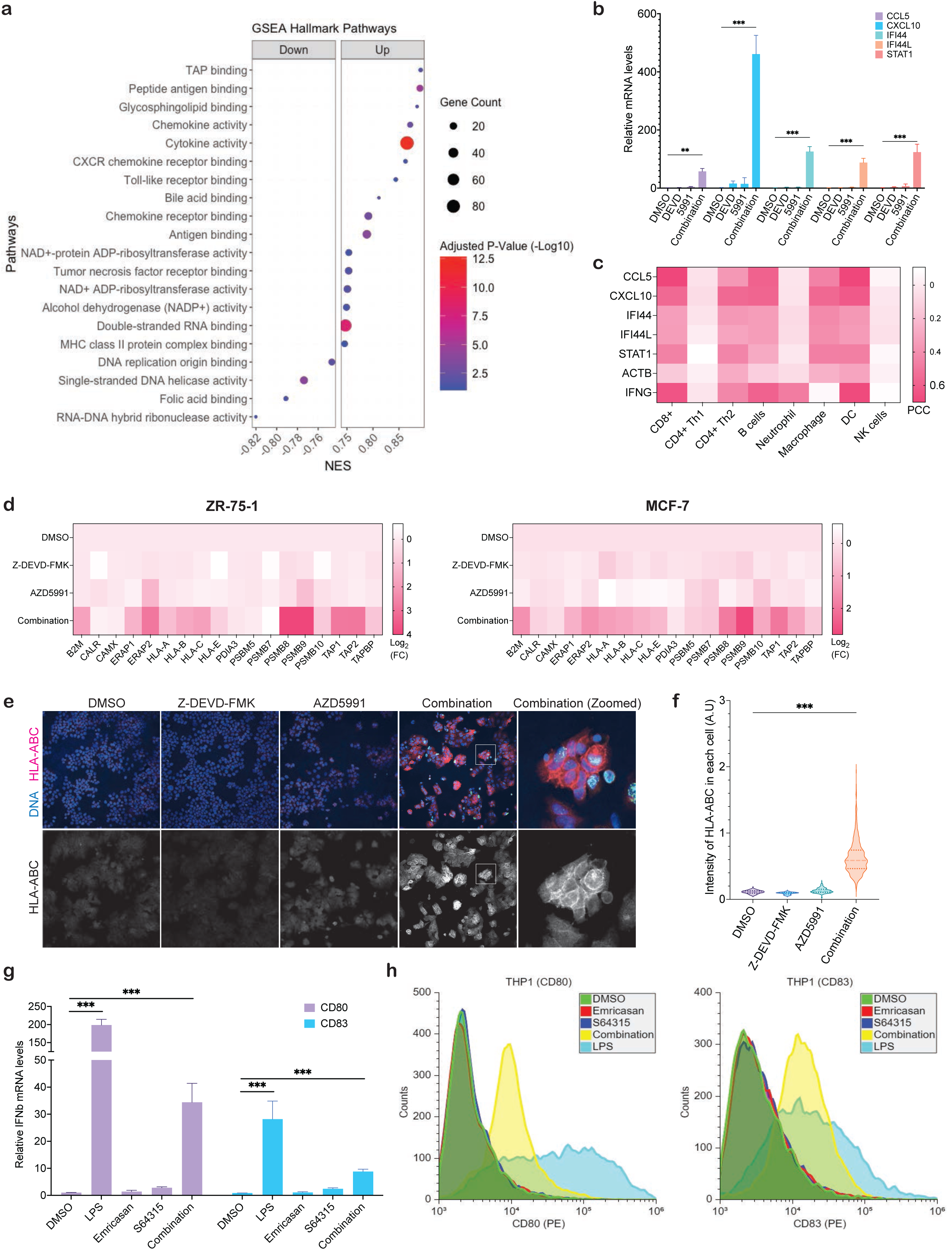
Enhancement of immune recognition markers and cytokine expression by combined MCL1 and caspase inhibition in breast cancer cells. **a,** Top 20 hallmark pathways in GSEA by iDEP DEGs ^89^ (DMSO vs combination treatment, n=2 biologically independent samples for each treatment condition). **b,** Quantitative PCR analysis of cytokines involved in immune cell recruitment (CCL5, CXCL10, IFI44, IFI44L, and STAT1). Data represent mean +/- SEM; unpaired t-test, n=3 biologically independent samples. **c,** Heatmap correlating the expression of the upregulated cytokines with immune cell infiltrates, including CD8+ T, B, and dendritic cells, in breast cancer samples from the TCGA dataset analyzed (n=1084 breast cancer tissues) with the xCell algorithm ^37^. PCC, the Pearson correlation coefficient. **d,** Heatmap displaying the transcriptional upregulation of antigen processing and presentation (APM) genes in ZR-75-1 (left) and MCF-7 (right) cells analyzed by qPCR. (n=3 biologically independent samples) **e,** Immunofluorescence (IF) analysis of expression of HLA-ABC in ZR-75-1 cells treated with a combination of MCL1 inhibitor AZD5991 (2µM) and caspase-3 inhibitor Z-DEVD-FMK (10 µM) versus MCL1 inhibitor alone. (n=3 biologically independent samples) **f,** Quantification of (e) by CellProfiler (Version 4.2.6). Data represent mean +/- SEM; unpaired t-test; n=3 biologically independent samples; 5 fields for each sample were analyzed. **g,** Quantitative PCR to measure expression of CD80 and CD83 in THP1 upon 24 hr incubation with the conditioned media obtained from ZR-75-1 treated with MCL1 inhibitor S64315 (2 µM) and caspase-3 inhibitor emricasan (10 µM). Data represent mean +/- SEM; unpaired t-test; n=3 biologically independent samples. **h**, Representative flow cytometry histogram analyzing the expression of CD80 and CD83 in THP1 upon 24 hr incubation with the conditioned media obtained from ZR-75-1 treated with MCL1 inhibitor S64315 (2 µM) and caspase-3 inhibitor emricasan (10 µM). GSEA, Gene Set Enrichment Analysis; DEGs, differentially expressed genes; NES, normalized enrichment score. Statistical significance is noted as *p < 0.05, **p < 0.01, ***p < 0.001.

We sought to examine how robust and acute activation of type I IFN response might impact immunogenicity. We confirmed that cytokines (CCL5, CXCL10, IFI44, IFI44L and STAT1) involved in the recruitment of immune cells ^36^ were highly upregulated by the combination treatment (Fig. 2b). Analysis of the association of these upregulated cytokines with immune cell infiltrates using TCGA dataset and xCell algorithm ^37^ demonstrated a strong association of cytokine expression with CD8+ T, B and dendritic cell (DC) infiltrates in over a thousand breast cancer samples (Fig. 2c). Furthermore, in ZR-75-1 and MCF-7 cells, combinatorial treatment upregulated transcription of most genes responsible for antigen processing and presentation (APM) ^38, 39^, a multistep process consisting of antigen peptide generation and loading of MHC class I molecules (Fig. 2d), suggesting an improved capability of tumor cells to process and present antigens ^40, 41^.

Because reduced expression of MHC class I molecules on the cell surface is one of the most important immune escape mechanisms in tumors ^42^ including breast cancer ^43–45^, we examined whether the combinatorial treatment would overcome this by enhancing expression of MHC class I molecules along with type I IFN response induction. Quantitative immunofluorescence (IF) was utilized to measure the expression levels of HLA-ABC proteins in ZR-75-1 cells. The results showed a significant increase in HLA-ABC expression when cells were treated with both MCL1 and caspase inhibitors, compared to treatment with MCL1 inhibitors alone (Fig. 2e, f). This suggests that the combination treatment can potentially restore immune recognition by increasing the visibility of cancer cells to the immune system.

To evaluate the potential impact of the combination treatment on immunogenicity by DC maturation in the tumor microenvironment, we examined the maturation of THP1-derived immature DCs (iDC) ^46, 47^ upon exposure to lipopolysaccharide (LPS) as a positive control and conditioned media from ZR-75-1 cells treated with the combination of emricasan and S64315. The conditioned media obtained from ZR-75-1 treated with the combination demonstrated increase in both mRNA levels and surface expression of DC maturation markers ^48^, CD80 and CD83 (Fig. 2g, h).

### The combination treatment triggers mitochondrial permeabilization leading to the release of mitochondrial DNA into the cytoplasm

Given the established role of MCL1 inhibitors in promoting MOMP ^49, 50^, we posited that such induction would lead to the cytosolic translocation of mitochondrial DNA (mtDNA). First, mitochondrial permeability was confirmed using the well-established method of quenching mitochondrial calcein AM fluorescence by cobalt upon mitochondrial permeability transition ^51^. In this assay, increased mitochondrial permeability is indicated by a reduction in non-quenched calcein AM fluorescence. In ZR-75-1 cells, MCL1 inhibition led to higher mitochondrial permeability, as evidenced by decreased calcein AM fluorescence intensity, with the remaining signal co-localizing with the mitochondrial marker Mitotracker (Fig. 3a, b and Extended Data Fig. 3). Second, we selectively permeabilized plasma membranes without disrupting other membrane-bound compartments including nucleus, mitochondria and endoplasmic reticulum to stain only cytoplasmic dsDNA. We confirmed that low concentration of saponin (below 0.005%) resulted in selective permeabilization of plasma membrane as previously reported (Extended Data Fig. 4a) ^52^. We performed quantitative analysis of cytoplasmic dsDNA foci (Extended Data Fig. 4b) and observed a significant increase in cytoplasmic dsDNA following MCL1 inhibition (Fig. 3c, d).

**Figure 3.**
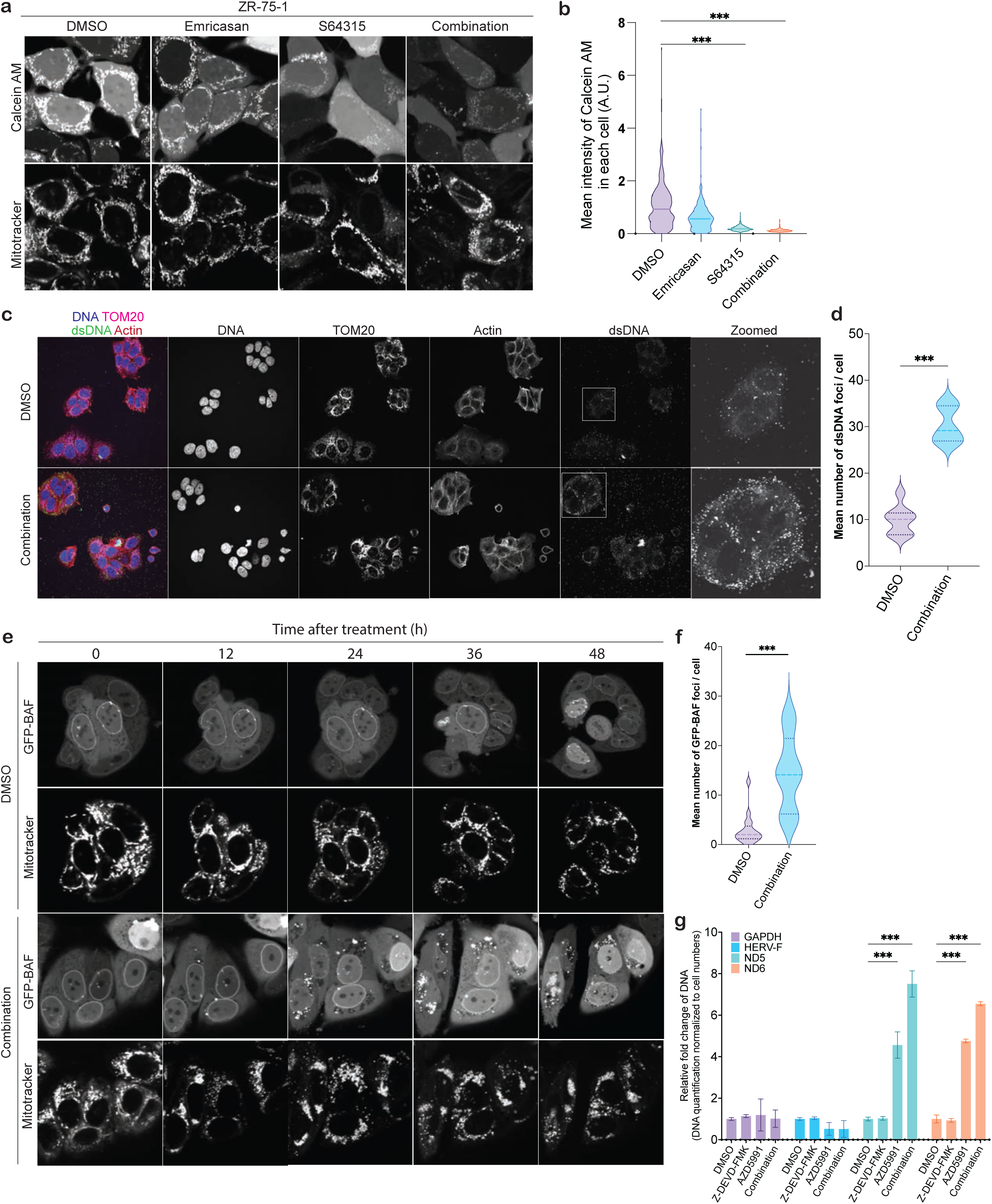
Release of mtDNA into cytoplasm by MCL1 and caspase inhibition. **a,** Representative live-cell imaging of WT ZR-75-1 cells showing calcein AM and MitoTracker™ Red FM. Cells were treated with a combination of MCL1 inhibitor S64315 (2 µM) and pan-caspase inhibitor Emricasan (10 µM). **b**, Quantification of (a) by CellProfiler (Version 4.2.6). **c**, Representative immunofluorescence (IF) staining (n=3 biologically independent samples) for cytoplasmic dsDNA (green), mitochondrial marker TOM20 (magenta), and actin (red) in MCF7 cells. **d,** Quantification of (c) by CellProfiler (Version 4.2.6). Data represent mean +/- SEM; unpaired t-test; n=3 biologically independent samples. **e**, Representative live-cell images (n=3 biologically independent samples) of ZR-75-1 expressing GFP-BAF treated with DMSO or combination in a time series with 12-hour intervals. **f,** Quantification of (e) 48 hr after treatment by CellProfiler (Version 4.2.6). Data represent mean +/- SEM; unpaired t-test; n=3 biologically independent samples. **g,** Quantitative PCR analysis of mitochondrial DNA (mtDNA) genes ND5 and ND6 in cytoplasmic fractions of ZR-75-1 upon the treatment with MCL1 inhibitor AZD5991 (2 µM) and/or caspase-3 inhibitor Z-DEVD-FMK (10 µM). Data represent mean +/- SEM; unpaired t-test; n=3 biologically independent samples.

To further confirm cytoplasmic dsDNA release, we generated ZR-75-1 cells stably expressing BAF1-GFP, a known marker for cytoplasmic dsDNA binding ^53^. The formation of BAF1 foci is indicative of cytoplasmic dsDNA release, as BAF1 binds to phosphate backbone of DNA to protect it from degradation ^54^. This approach enabled the detection of cytoplasmic DNA under live-cell imaging conditions for 72 hours. The live-cell imaging revealed a significant increase in the formation of BAF1 foci in cells treated with the combination therapy, compared to controls (Fig 3.e, f and Supplementary Movies 1 and 2). In addition to BAF1 foci formation, mitochondrial clustering around the nucleus, a phenotype often associated with mitochondrial stress ^55–57^, was observed in cells subjected to the combination treatment.

Lastly, to perform quantitative analysis of cytoplasmic mtDNA, we first fractionated cell lysates and confirmed the purity of fractionation by immunoblot (Extended Data Fig. 5a). Mitochondrial genes ND5 and ND6 ^58^ (surrogate markers of mtDNA) were confirmed by qPCR to be present exclusively in the mitochondrial fraction and whole-cell lysate in unstimulated cells (Extended Data Fig. 5b). In contrast, analysis of cytoplasmic fractions revealed markedly elevated ND5 and ND6 levels following treatment with MCL1 inhibitor alone or in combination with a caspase inhibitor without a significant effect on the levels of GAPDH and endogenous retroviral gene HERV-F (Fig. 3g). These results indicate that while MCL1 inhibition alone can initiate DAMP release, a robust type I IFN response requires concurrent caspase inhibition, suggesting a synergistic effect on the immune signaling cascade. Altogether, these findings strongly support the conclusion that mitochondrial permeabilization induced by the combination treatment facilitates the release of mitochondrial DNA, thereby amplifying immune-related pathways.

### The IFN response caused by the combination treatment requires STING

To elucidate the pathway responsible for IFN induction triggered by the combination treatment, we utilized isogenic MCF7 cell lines with CRISPR/Cas9-mediated knockout (KO) of key pattern recognition receptors (PRRs), including cGAS, STING, RIG-I, MDA5, and MAVS (Fig. 4a). These PRRs are integral to the detection of cytoplasmic DNA or RNA and subsequent activation of innate immune responses. Successful gene knockouts were confirmed by immunoblotting (MDA5 expression upon poly I:C treatment) (Fig. 4b). The cytotoxic effects of the combination treatment were comparable across all PRR-KO cell lines and Cas9 control cells, indicating that any observed differences in IFN response could be attributed specifically to the absence of the targeted PRR genes rather than to variations in cytotoxicity (Extended Data Fig. 6a).

**Figure 4.**
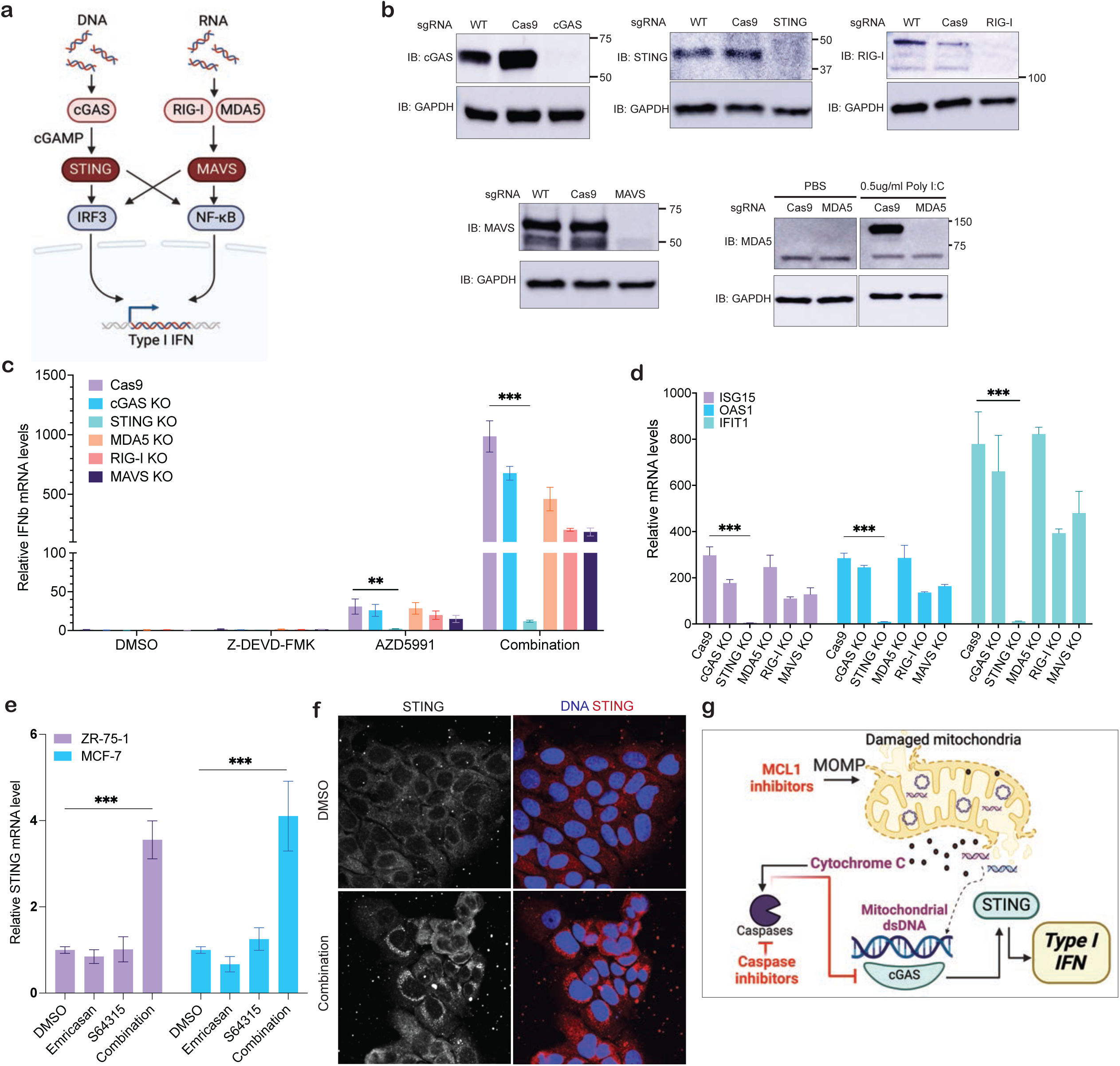
The STING-dependent IFN response induced by MCL1 and caspase inhibition. **a,** The pattern-recognition receptor pathway recognizing either cytoplasmic DNA or RNA and inducing type I IFN response. **b,** Western blot of CRISPR/Cas9-mediated knockout (KO) in MCF7 cell lines targeting cGAS, STING, RIG-I, MDA5, and MAVS. (n=3 biologically independent samples) **c,** Quantitative PCR analysis of IFNB1 expression in the knockout cell lines following treatment with DMSO, 10 µM Z-DEVD-FMK (caspase-3 inhibitor), 2 µM AZD5991 (MCL1 inhibitor), and their combination. **d,** Quantitative PCR analysis of IFN-stimulated genes (ISGs) in the KO cell lines following treatment with DMSO, 10 µM Z-DEVD-FMK (caspase-3 inhibitor), 2 µM AZD5991 (MCL1 inhibitor), and their combination. **e**, Quantitative PCR analysis of STING in ZR-75-1 and MCF-7 following treatment with DMSO, 10 µM emricasan, 2 µM S64315 (MCL1 inhibitor), and their combination. **f**, Representative immunofluorescence (IF) staining (n=3 biologically independent samples) for STING in ZR-75-1 cells treated with DMSO or combination of 10 µM Z-DEVD-FMK (caspase-3 inhibitor) and 2 µM AZD5991 (MCL1 inhibitor). **g**, Suggested model for the current study (Created with BioRender.com). Data represent mean +/- SEM; unpaired t-test; n=3 biologically independent samples. Significance is noted as *p < 0.05, **p < 0.01, ***p < 0.001. **f**, Suggested model for this study.

While knockouts of most PRRs resulted in a partial reduction in IFNB1 expression, the absence of STING led to a complete abrogation of IFNB1 induction and other ISGs, including ISG15, OAS1, and IFIT1 (Fig. 4c, d). This result underscores STING’s pivotal role as the central mediator of the type I IFN response in this system. Additionally, previous studies have reported transcriptional regulation of STING and a positive feedback loop in the IFN response ^59, 60^. Therefore, we examined the mRNA levels of STING in ZR-75-1 and MCF-7 cells following treatment. Consistent with the literature, we observed upregulated STING transcriptional expression, suggesting that this positive feedback loop enhances type I IFN signaling pathway (Fig. 4e). To further confirm STING involvement, we investigated its perinuclear clustering, a hallmark of STING activation, which involves its translocation and clustering around the perinuclear region to initiate downstream signaling events ^61, 62^. Indeed, perinuclear clustering of STING was exclusively and distinctly observed following the combination treatment in ZR-75-1 cells (Fig. 4f).

Together, these results demonstrate that STING activation is essential for driving the observed type I interferon response, as evident by the increased expression levels of downstream ISGs, enhanced transcriptional activity of STING itself, and the complete abrogation of the IFN response in STING knockout cells (Fig. 4g).

### The combination treatment inhibits tumor growth *in vivo* in IFN-dependent manner

We next proceeded to test the effect of the combination of MCL1 and caspase inhibition in an immunocompetent syngeneic murine model of breast cancer. Because MCL1 inhibitors have been reported to exhibit about 6-fold lower specificity in mouse cells^63^, we first assessed IFN response in murine syngeneic breast cancer lines, including 67NR, 4T1, MC7-L1^64^, and MC4-L2 ^64^, to select an optimum cell line for our *in vivo* study. Because MC4-L2 showed the most robust IFN response to the combination treatment, it was selected for further study (Extended Data Fig. 7a). To identify the most effective combination of MCL1 and caspase inhibitors in MC4-L2, we tested several MCL1 inhibitors including S63845 (Servier), S64315 (Servier), A1210477 (Abbvie) and AZD5991 (AstraZeneca) in combination with the pan-caspase inhibitor emricasan, which has been shown to have a favorable safety profile in early phase clinical trials ^65, 66^. Among these, the combination of S64315 and emricasan elicited the strongest IFN response (Extended Data Fig. 7b). This was further confirmed by measuring IFN-β secretion using ELISA (Extended Data Fig. 7c). Consistent with observations in human breast cancer cell lines, we also confirmed a significant increase in cytoplasmic mtDNA (ND5 and ND6) release in MC4-L2 cells following the combination treatment, as demonstrated by cell fractionation and PCR analysis (Extended Data Fig. 7d).

For the *in vivo* study, MC4-L2 cells were injected orthotopically into the mammary fat pads of immunocompetent BALB/C mice. Tumor growth was monitored, and once the tumors reached a volume of 200 mm^3^ (day 0), treatment was initiated. The mice were administered S64315 at a dose of 40 mg/kg (i.v.) and/or emricasan at a dose of 10 mg/kg (i.p.) continuously for 5 days (Fig. 5a and Extended Data Fig. 7e). There was no significant change in the body weight of mice in all treatment groups during the drug treatment period, indicating no substantial toxicity from the combination treatment (Extended Data Fig. 7f). Strikingly, the combination treatment exhibited significant inhibitory effect on tumor growth to a much higher degree than either MCL1 inhibitor or caspase inhibitor alone (Fig. 5b). To examine whether this effect was associated with an increase in IFN levels, we measured serum IFN-β levels collected prior to drug administration and at days 3 and 6 following initiation of drug treatment. Indeed, we observed a significant increase in serum IFN-β levels in the combination treatment group at day 6 (Fig. 5c). To determine if downstream ISGs are similarly induced by the combination *in vivo*, we performed RNA-seq on vehicle and combination treatment tumor samples from mice sacrificed on day 15. We employed GSEA to query the differentially expressed genes against the “Hallmark” database (Fig. 5d) and GO gene set (Extended Data Fig. 7g). Consistent with the hypothesis that the combination treatment correlates with tumor immunogenicity and upregulation of the antigen presentation pathway, both pathway analyses demonstrated robust enrichment in cytokine signaling and MHC protein complex gene expression, and concomitant upregulation of IFN response genes.

**Figure 5.**
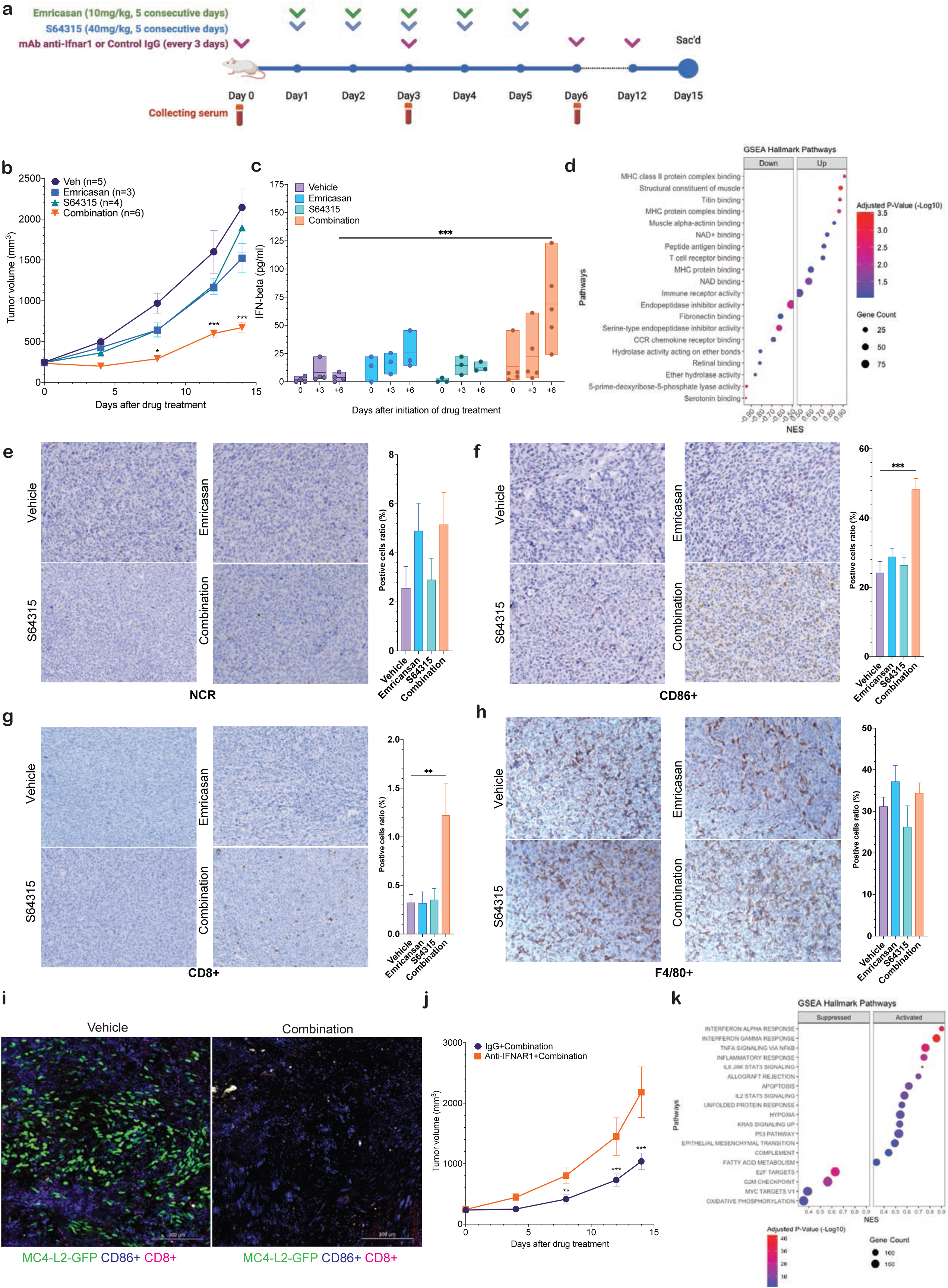
*In vivo* assessment of MCL1 and caspase inhibitor combination treatment and its impact on the tumor microenvironment. **a,** Treatment timeline illustrates the administration of S64315 (40mg/kg) and emricasan (10mg/kg) over five consecutive days, with serum collection and anti-IFNAR1 or control IgG antibody treatment at indicated intervals (Created with BioRender.com). **b,** Murine breast cancer cell line (MC4-L2) was injected into the mammary fat pad of wildtype mice (BALB/c) treated with S64315 and/or emricasan. Tumor growth curves of the progression of MC4-L2 tumors in mice (Veh, n=5; emricasan, n=4; S64315, n=6; Combination, n=8). Data represent mean +/- SEM; two-way ANOVA. **c,** IFN-β concentration in serum, measured by ELISA before and after treatment, indicates an increase in the combination treatment group. Data represent mean ± SEM; one-way ANOVA, Veh, n=5; emricasan, n=4; S64315, n=6; Combination, n=8 **d,** Top 20 hallmark pathways in GSEA by iDEP DEGs (Vehicle vs combination treatment, n=3 mice for each treatment condition). **e, f, g and h,** Immunohistochemistry (IHC) analysis and quantification for immune cell markers NCR (NK cells) (e), CD86 (dendritic cells) (f), CD8 (CD8+ T cells) (g) and F4/80 (macrophages) (h), in tumor sections. Quantification of positive stained cells per field from treated tumors (Veh, n=5; emricasan, n=4; S64315, n=6; Combination, n=8). Quantification data represent mean ± SEM; unpaired t-test; 3-5 images for one IHC slide. **i,** MC4-L2 was injected into the mammary fat pad of wildtype mice (BALB/c) treated with S64315 and emricasan. Prior to these treatments, mice were pre-treated with either IgG or anti-IFNAR1 monoclonal antibody. Tumor growth curves of the progression of MC4-L2 tumors in mice (IgG+combination, n=7; anti-IFNAR1+combination, n=8). **j**, Top 20 hallmark pathways in GSEA by iDEP DEGs (anti-IFNAR1 vs control IgG, n=5 mice for each treatment condition). **k**, Infiltration level of CD8+ T cell and mDC in *in vivo* RNAseq data analyzed by timer.cistrome.org webtool. Data represent mean +/- SEM; unpaired t-test, n=4 biologically independent samples. Data represent mean +/- SEM; two-way ANOVA. Statistical significance is noted as *p < 0.05, **p < 0.01, ***p < 0.001 and ns, statistically not significant.

**Figure 6.**
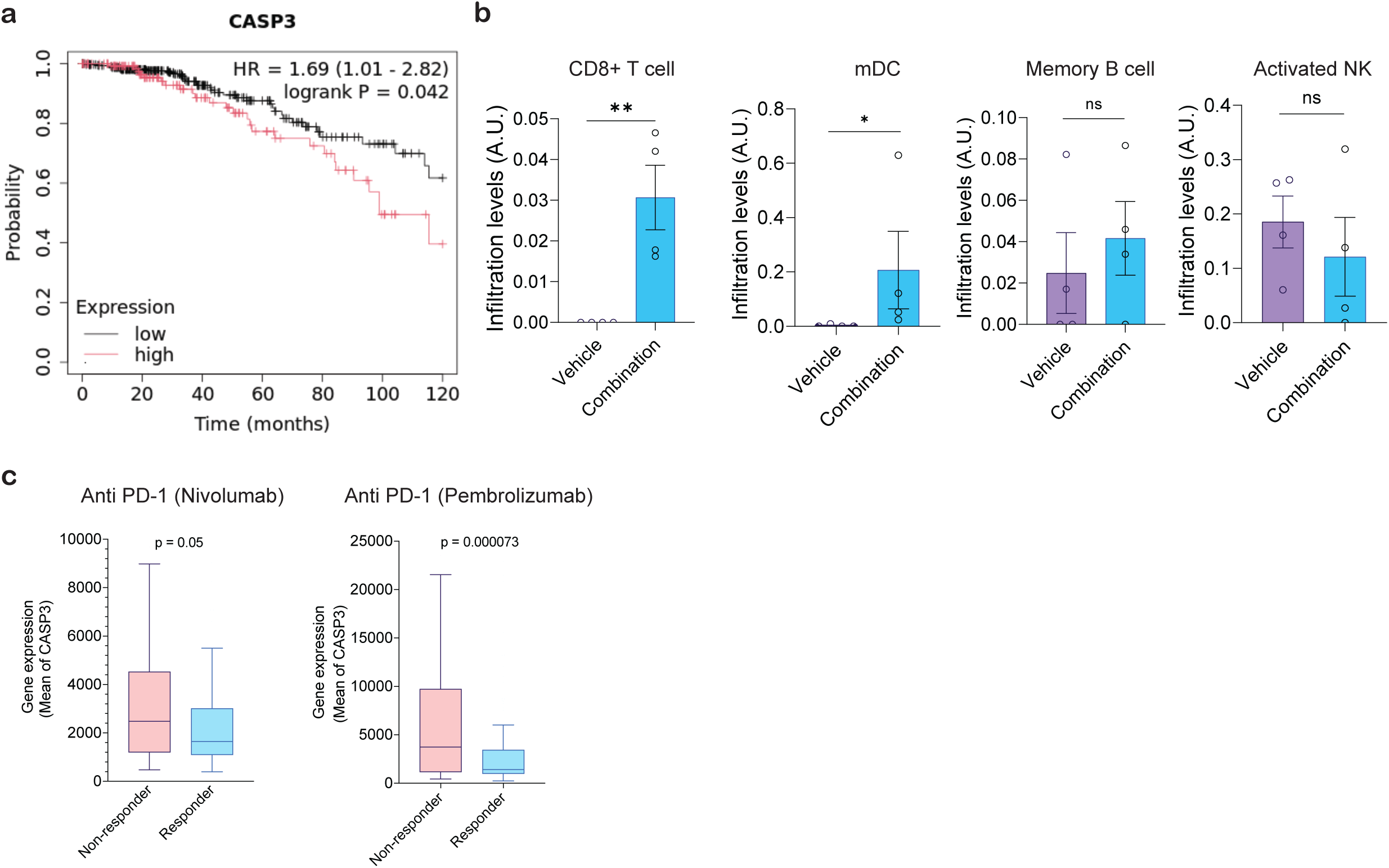

We next investigated the impact of combination treatment on tumor microenvironment by immunohistochemistry (IHC). While there was no significant change in the number of NCR+ (NK cells) and F4/80+ cells (macrophages) (Fig. 5e and 5h), our analysis revealed an increase in CD86+ dendritic cells and CD8+ T cells in tumors treated with the combination treatment (Fig. 5f and 5g) consistent with activation of anti-tumor immune response.

To further assess the impact of the combination treatment on tumor dynamics and visualize the tumor cell response within the treated microenvironment, we performed *in vivo* imaging using MC4-L2-GFP cells immediately after completing 5 consecutive days of treatment. The imaging results showed a striking reduction in tumor cell fluorescence in the combination treatment group, indicating a significant decrease in tumor burden compared to the vehicle group (Extended Data Fig. 8a, 8b, and 8c). To confirm gene expression changes in the tumor microenvironment, we performed RNA-seq on tumor tissues after the 5-day treatment (Extended Data Fig. 8d). This revealed striking increases in genes related to cytolysis and leukocyte-mediated cytotoxicity, further highlighting the strong immune response and anti-tumor effects induced by the combination treatment.

To determine whether the observed tumor growth inhibition by the combination treatment was due to immune-mediated effects rather than direct cytotoxic or cytostatic effects of the drugs, we administered anti-interferon-α receptor 1 (IFNAR1) neutralizing monoclonal antibodies (mAb) to mice before and throughout the drug treatment while monitoring tumor growth. Importantly, introduction of an anti-IFNAR1 mAb abrogated the tumor-suppressive effect of the combination treatment (Fig. 5i), demonstrating that anti-tumor effects of the combination treatment are dependent on IFN pathway. Notably, no significant toxicity was observed in the IFNAR1-blocking group, consistent with prior treatment conditions (Extended Data Fig. 9a). Moreover, RNA-seq analysis confirmed that only mice treated with the control IgG antibody but not IFNAR1 antibody exhibited an increase in cytokine, antigen presentation, and IFN response-related genes (Fig. 5j and Extended Data Fig. 9b and 9c), reinforcing the conclusion that the anti-tumor activity of the combination treatment is dependent on type I IFN response induction by the treatment.

### Caspase-3 inhibits innate immune signaling and shapes immunotherapy resistance and poor survival in breast cancer

Caspase-3, classically known for its role in executing apoptosis, has recently been identified as a key suppressor of innate immunity through the direct cleavage of multiple PRRs, including cGAS, MAVS, and IRF3 ^26, 67^. Although caspase-3 shares some substrate specificity with other executioner caspases ^68^, it is the only caspase demonstrated to directly cleave PRRs such as cGAS and MAVS, thereby uniquely acting as a direct suppressor of type I IFN signaling.

To assess the clinical significance of caspase-3 expression in breast cancer, we analyzed datasets from The Cancer Genome Atlas (TCGA) and The Genotype-Tissue Expression (GTEx). CASP3 expression was significantly elevated in breast cancer tissues compared to normal controls (Extended Data fig.10a). Because immunosuppression can be associated with a worse prognosis, we analyzed whether the expression of caspase-3 in breast cancer tissues with low mutation burden correlates with overall survival (https://kmplot.com/) ^69^. The analysis revealed a statistically significant negative correlation between overall survival and the expression of CASP3 with HR of 1.69 (Extended Data fig.10b). Furthermore, analysis of immune infiltration signatures revealed a significant inverse correlation between CASP3 expression and the abundance of immune effector cells, including CD8+ T cells, memory B cells, dendritic cells, and activated NK cells (Extended Data Fig. 10c) (http://timer.cistrome.org) ^69^. To corroborate these findings, we assessed immune cell infiltration in tumor tissues obtained from our *in vivo* experiments. The transcriptome analysis revealed increased infiltration of CD8+ T cells and mDCs in the combination treatment group (Fig. 5k and Extended Data Fig. 10d), aligning with the IHC results (Fig. 5e-h). This suggests that caspase-3 may not only suppress innate immune signaling through PRR cleavage, but also actively shape more immunosuppressive microenvironment.

Consistent with this, analysis of over 1,400 tumor samples from diverse cancer types demonstrated that patients with lower CASP3 expression exhibited significantly improved responses to anti-PD-1 therapies such as nivolumab and pembrolizumab (Extended Data Fig. 10e). Together, these findings highlight caspase-3 as a potential negative regulator of tumor immunogenicity and a candidate prognostic and predictive biomarker for immunotherapy outcomes.

## Discussion

Our study introduces a novel strategy to treat immunologically “cold” cancers, including ER+ breast cancer ^70, 71^, using a combination of MCL1 and caspase inhibition. By producing a robust and acute type I IFN response specifically in tumor cells, this strategy has an important advantage over existing immune-based strategies such as STING agonists ^72^. In addition, the study uncovers the important immunomodulary role of MCL1 inhibitors, which have been successfully used in the treatment of hematologic malignancies and are being tested in several epithelial cancers including breast adenocarcinoma ^73^. Importantly, both MCL1 and caspase inhibitors have been shown to be safe in early phase clinical trials ^66, 74^, allowing for a more rapid translation of this treatment strategy to patients.

The combination treatment effectively suppressed tumor growth in immunocompetent syngeneic mouse models, reprogramming the tumor microenvironment to support a more robust anti-tumor immune response. Although the combination did not fully eradicate tumors in mice, we anticipate a more pronounced therapeutic impact in human tumors for several reasons. First, MCL1 inhibitors have been shown to display roughly six-fold higher specificity in human cells compared to mouse cells ^63^. Second, type I IFN response can induce PD-L1 expression ^75^, and our analyses reveal that low CASP3 expression correlates with favorable outcomes following anti-PD-1 therapy. Consequently, combining dual MCL1 and caspase inhibition with immune checkpoint blockade could further enhance therapeutic efficacy, a strategy that will be pursued in future investigations.

It can be argued that caspase inhibition is counterintuitive as a cancer strategy as it can suppress apoptotic cell death. Importantly, we have demonstrated that caspase inhibition has no significant effect on cytotoxicity in cancer cell lines. Furthermore, unlike hematologic malignancies, most solid cancers including carcinomas and sarcomas are resistant to apoptotic cell death and are believed to die primarily by other mechanisms ^76–79^. This strategy is thus particularly attractive for tumors that are resistant to conventional cytotoxic treatment strategies. Dual MCL1 and caspase inhibition offers several important advantages over existing strategies aimed at inducing type I IFN response. First, even though the drugs are systemically administered, type I IFN response appears to be limited to cancer cells, thus reducing the likelihood of acute toxicity. The favorable safety profile of systemic administration of both MCL1 and caspase inhibitors from early phase clinical trials (Table 1) ^65, 73, 80^ offers a significant advantage over other strategies to induce cGAS-STING activation such as STING agonists, which suffer from a potential adverse effect of excessive cytokine induction upon systemic administration ^81^. This largely limits STING agonist administration to intratumoral injection, which has limitations including tumor accessibility and insufficient retention ^81^.

**Table 1.**
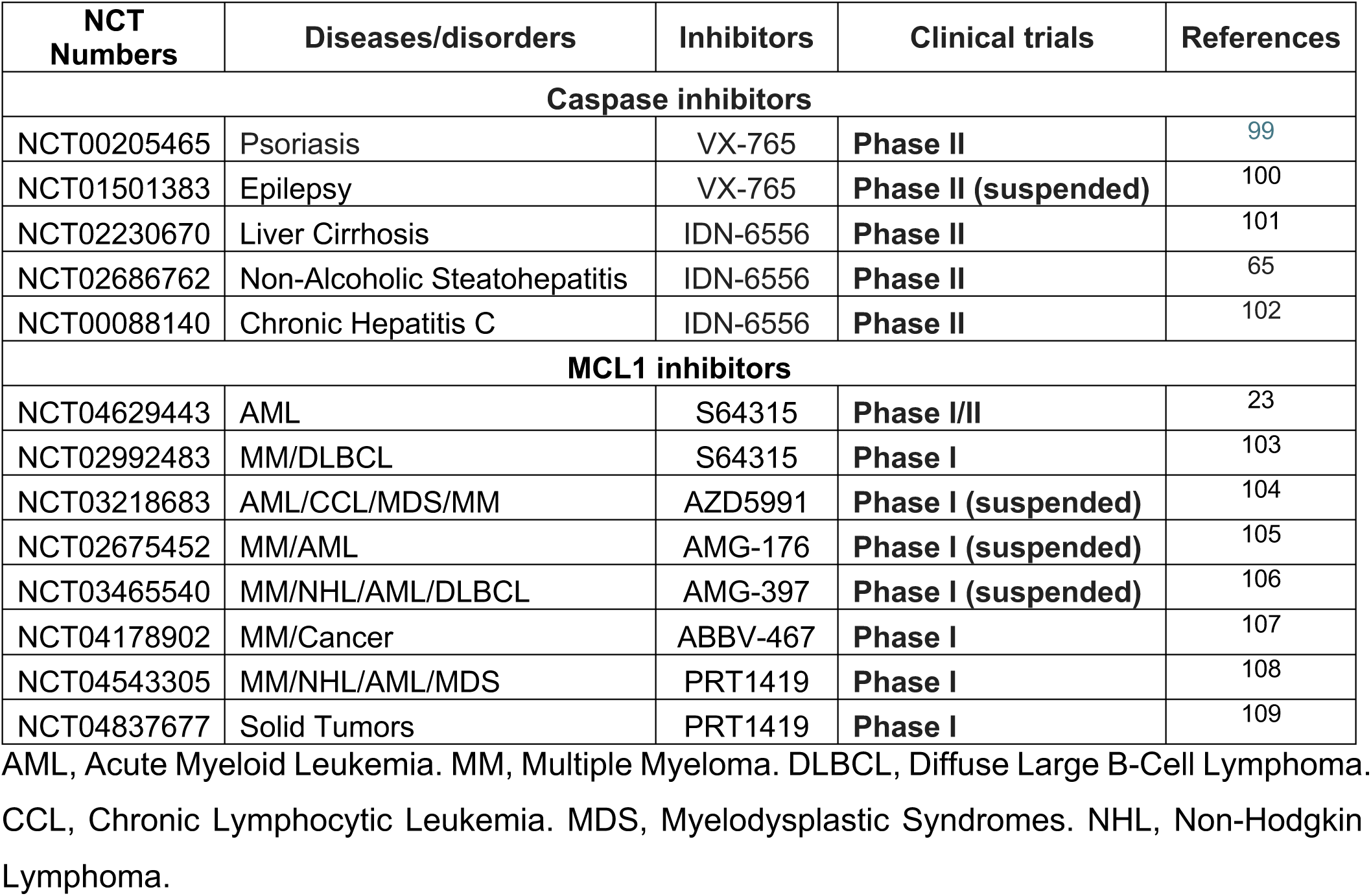
Clinical trials of caspase or MCL1 inhibitors.

In conclusion, our study proposes a drug combination treatment that can enhance anti-tumor immunity with the potential for rapid clinical translation in breast cancers and other malignancies. This strategy can be effectively combined with immune checkpoint inhibitors to further enhance anti-tumor immune response. Furthermore, by upregulating antigen presentation machinery specifically in tumor cells, this strategy has the potential to synergize with efforts to target tumor-specific peptide-MHC epitopes using radiopharmaceuticals, bi-specific T cell-engagers (BITEs) and CART-T cells ^82–85^.

## Online Methods

### Cell lines and culture conditions

Human breast cancer cell lines MCF-7, ZR-75-1, and T47D were cultured in Dulbecco’s Modified Eagle Medium (DMEM) from Gibco, enriched with 10% fetal bovine serum (FBS), 100 U/mL penicillin, and streptomycin. Murine breast cancer cell lines 4T1, MC7-L1, and MC4-L2 were grown in RPMI 1640 medium (Gibco) supplemented with 10% FBS, 100 U/mL penicillin, and streptomycin. Non-transformed cell lines BJ-hTERT and RPE1-hTERT were maintained in DMEM (Gibco) and DMEM/F-12 (Gibco), respectively, each supplemented with 10% FBS, 100 U/mL penicillin, and streptomycin. We conducted regular checks for mycoplasma contamination and authenticated cell lines using the MycoAlert Mycoplasma Detection Kit and molecular diagnostics. Murine breast cancer cell lines MC7-L1 and MC4-L2 were kindly provided by Richard Schwartz, Ph.D. (Michigan State University). PRRs including cGAS, STING, MAVS, RIG-I and MDA5 KO MCF7 cells were kindly provided by Qin Yan, Ph.D. (Yale University). Stable MC4-L2-GFP cells or ZR-75-1 expressing GFP-BAF1 and RFP-H2b were generated by transduction using lentivirus. In brief, to produce lentiviral particles, HEK 293 cells were transfected with 12 µg lentiviral plasmid carrying GFP-H2B, RFP-H2B or GFP-BAF1 (Spektor laboratory) ^86^ + 7.5 µg pMD2.G (#12259 AddGene) + 6 µg psPAX2 (#12260, AddGene). Then, cells were infected for 24 h with viral particles in the presence of 10 μg/ml polybrene, washed, and allowed for 48 h to recover before fluorescence-activated cell sorting.

### CRISPR/Cas9 knockout generation

PRRs KO MCF-7 cell lines (cGAS, STING, MDA5, RIG-I and MAVS), a kind gift of Dr. Yan (Yale University), were subjected to single-cell sorting for GFP positive cells by the Dana-Farber Cancer Institute Flow Cytometry Core and western blot analysis to verify KO in MCF7 cells.

### Animal experiments

For each mouse, 2.5×10^6^ MC4-L2 cells were prepared for each mouse and resuspended in 50 μl of PBS. Cells were injected into the fourth mammary fat pad of 8-week-old BALB/C (Charles River Laboratories) mice. Treatment was initiated when tumors reached approximately 200 mm3 (Tumor formula, 0.5 x length x width2). Both emricasan (MedChem Express, 10 mg kg-1 i.p.) and S64315 (MedChem Express, 40 mg kg-1 i.v.) were diluted with DMSO with 5% DMSO, 40% PEG300, 5% Tween80 and 50% DW and administered for 5 consecutive days. Anti-IFNAR1 antibody (200 ug per mouse, i.p.) and mouse IgG1 isotype control (200 ug per mouse, i.p.) were diluted in PBS and intraperitoneally administered every 3 days. Vehicle, emricasan, S64315, and anti-IFNAR1 antibody+emricasan+S64315 treatment groups were sacrificed on day 14. Emricasan+S64315 and IgG1+emricasan+S64315 treatment groups were sacrificed on day 22. Tumors were excised and portions were snap-frozen for RNA analysis, fixed in 10% buffered formalin for IHC. All studies were performed according to protocols approved by the UMass Medical School Institutional Animal Care and Use Committee.

### Western Blot

The cells were collected and disrupted in a solution composed of 150 mM NaCl, 20 mM HCl (pH 7.5), 0.5% NP-40, 5mM EDTA, and 5% glycerol, supplemented with protease and phosphatase inhibitors from Roche. Lysis was performed for 30 minutes on ice. The protein content was then quantified utilizing the BCA Protein Assay (#A55860, ThermoFisher Scientific). For electrophoresis, 20 or 50 μg of the lysate was mixed with 4X SDS loading buffer (#1610747, BioRad) and then loaded onto 4%-12% precast gels (NP0335, Life Technologies). Subsequent to the electrophoretic separation, proteins were transferred onto membranes via iBlot 2 Gel Transfer Device (Invitrogen). To prevent non-specific binding, the membranes were then incubated with 5% milk in TBST for 1 h at 24 °C. Primary antibody incubation (details in Reporting Summary) was performed overnight in a 4 °C cold room. This was followed by the application of suitable enhanced chemiluminescence secondary antibodies (also detailed in Reporting Summary) for 1 h at 24 °C. The membranes were developed using ECL Western Blotting Substrate (#32109, ThermoFisher Scientific) and visualized on an AI600 Chemiluminescent Imager (Amersham).

### Immunohistochemistry

Fixed tissues were paraffin-embedded and immunohistochemical staining was performed by iHisto. Stained tumor sections were viewed on an Olympus BX41 light microscope (Olympus). Images were captured with an Evolution MPColor camera (Media Cybernetics).

### Immunofluorescence

Cells were plated onto coverslips, and 2 or 3 days after drug treatment, they were fixed in 4% formaldehyde for a duration of 20 minutes at 24 °C. After fixation, the cells were permeabilized in 0.1% Triton X-100 for 5 minutes. For selective permeabilization of plasma membranes for dsDNA detection as previously described ^87^, cells were instead incubated in 0.005% saponin for 5 minutes. The coverslips were then rinsed three times with PBS and blocked with 3% BSA in PBS. Primary antibody incubation was performed (refer to Reporting Summary for antibodies and dilutions used) overnight in a 4°C cold room using a 3% BSA in PBS solution. Following primary antibody incubation, the coverslips were washed three times and incubated with the corresponding fluorescent secondary antibody in a 3% BSA in PBS solution for 1 h at 24 °C (refer to Reporting Summary for antibodies and dilutions used). Subsequently, the coverslips were washed three times with PBS and then incubated with Hoechst 33342 in PBS (1:2500 dilution, #62249, ThermoFisher Scientific) for 20 minutes at 24 °C. The coverslips were then mounted onto glass slides using ProLong™ Diamond Antifade Reagent (#P36961, ThermoFisher Scientific). Imaging was performed on a spinning disk confocal microscope (a Nikon Ti2 with a Yokogawa CSU-W1 spinning disk head) with a 40×/0.75 NA Plan Fluor Air or a 60×/1.42 NA Plan Apochromat oil immersion objective (Nikon). The images were analyzed and quantified using ImageJ/Fiji software (NIH) or CellProfiler (Broad Institute, Version 4.2.6).

### Fluorescence labeling and confocal live-cell microscopy

Cells were plated in iBidi µ-Plate 24 Well (#82426, iBidi) and then treated with 25nM MitoTracker™ Deep Red FM (#M22426, Invitrogen) for approximately 45 min-1 h before imaging. Cells were washed with PBS three times. Cells were imaged with or without drug treatments (2µM S64315 and 10µM emricasan). Imaging was performed on a spinning disk confocal microscope (a Nikon Ti2 with a Yokogawa CSU-W1 spinning disk head) with a 60×/1.42 NA Plan Apochromat oil immersion objective (Nikon). Laser excitation used: 488-nm, 561-nm, and 642-nm lasers. An Okolab CO2 incubator and controller were used to maintain samples at 37°C and 5% CO_2_. Time-lapse images were acquired at 15-min intervals and image acquisition was stopped 72 hours after the treatment. The images were analyzed and quantified using ImageJ/Fiji software (NIH) or CellProfiler (Broad Institute, Version 4.2.6).

### Differentiation of THP-1 cells into iDCs and incubation with conditioned media

To generate iDCs, 2 × 10^5^ THP-1 cells/mL were seeded in 5 mL of RPMI medium supplemented with 10% FBS, 1% PenStrep in a T25 flask. For differentiation, 100 ng/ml recombinant human GM-CSF and 100 ng/ml recombinant human IL-4 were added. The cells were incubated for 5 days at 37 °C with 5% CO_2_, with medium exchange and addition of fresh cytokines on day 3. For preparation of conditioned media, ZR-75-1 cells were treated with various drug combinations for 48 h, then washed three times with culture medium and three times with phosphate-buffered saline (PBS). Fresh medium was added, and the cells were incubated for an additional 48 h. The conditioned media were collected, centrifuged at 3000 rpm for 5 minutes, and filtered through a 0.22 µm filter before being used to treat iDCs. The iDCs were cultured with the conditioned media for 24 hours and then harvested for further analysis.

### Flow cytometry

Cells were stained with the indicated antibodies (see Reporting Summary) at 4 °C for 30 minutes and then fixed with 3% formaldehyde. Following this, the cells were washed, resuspended in PBS, and analyzed using a BD LSR Fortessa flow cytometer. Analysis was carried out using Floreada.io (https://floreada.io/)

### Cytoplasmic fractionation and cytoplasmic mtDNA quantification

Cell fractionation and mtDNA quantification were conducted based on the protocol described in Bryant et al. ^88^. Briefly, cells treated with different drugs were pelleted by centrifugation and lysed using Digitonin Lysis Buffer (50 mM HEPES pH 7.4, 150 mM NaCl, 18 μg/mL digitonin, protease inhibitors). The supernatant obtained, representing the cytosolic extract, was collected, with a portion set aside for western blotting. The pellet was then washed and lysed in NP-40 Lysis Buffer (50 mM Tris pH 7.5, 150 mM NaCl, 1 mM EDTA, 1% NP-40 (v/v), 10% glycerol (v/v), protease inhibitors), facilitating the release of mtDNA from mitochondria while preserving the nuclear integrity. This process resulted in the mitochondrial extract, with a part reserved for western blotting. The remaining nuclear pellet was processed using SDS Lysis Buffer (20 mM Tris pH 8, 1% SDS (v/v), protease inhibitors) and sonication to extract nuclear DNA, producing the nuclear extract, a portion of which was also set aside for Western blotting. DNA extraction from each fraction was performed using the PureLink™ Genomic DNA Mini Kit (#K182002, Invitrogen), and mtDNA quantification was conducted by qPCR with primer sets listed in Supplementary Table 1. Successful fractionation was confirmed by western blot using the following antibodies: anti-TOM20 for mitochondrial, anti-Histone H3 for nuclear, and anti-GAPDH for cytoplasmic fractions.

### Mitochondrial permeability assay (Calcein AM)

ZR-75-1 cells were plated in iBidi µ-Plate 24 Well (#82426, iBidi). Two days after drug treatment, cells were stained with 1 µM Calcein AM, Hoechst 33342 (1:2500 dilution, #62249, ThermoFisher Scientific) and 25nM Mitotracker Deep Red FM (#M22426, ThermoFisher Scientific) with 1 mM CoCl₂, a quencher of calcein AM. Cells were incubated with the combined staining solution for 60 minutes at 37°C. Live-cell imaging was performed using a Nikon Ti2 spinning disk confocal microscope (a Yokogawa CSU-W1 spinning disk head and a 60×/1.42 NA Plan Apochromat oil immersion objective). Mitochondrial permeability was assessed by monitoring the retention of calcein fluorescence within mitochondria in the presence of CoCl₂

### Real-time quantitative polymerase chain reaction (RT-qPCR)

Total RNA extraction was performed utilizing the Qiagen RNeasy kit (#74004). RNA concentration was measured using a Nanodrop spectrophotometer and then RNA was used to generate complementary DNA using the SuperScript III First-Strand Synthesis System (#18080400, Invitrogen) following the manufacturer’s protocol. RT-qPCR analyses were conducted on a StepOnePlus™ Real-Time PCR System (Applied Biosystems) with qPCR SuperMix (#10572014, ThermoFisher Scientific). For evaluating relative gene expression, the delta-delta Ct method (2^−ΔΔCt)^ was applied, using GAPDH as the normalization control. Details of the primer sequences are listed in Supplementary Table 1.

### IFN-β protein concentration assay

Cells seeded in 6-well plates were stimulated as demonstration in the figure legends. Supernatants were harvested, and the amount of secreted IFN-β protein was quantified with the Human Human IFN-β bioluminescent ELISA kit (invivogene) or Murine IFN-β bioluminescent ELISA kit (invivogene). To measure IFN-β protein concentration in mouse serum, blood was collected by submandibular puncture on days −1, +2, +5 and allowed to clot at 24 °C followed by centrifugation at 2000 g for 10 minutes, 4 °C. The resulting supernatants were then used for the serum analysis.

### RNA sequencing

Library preparation, sequencing, and subsequent data analysis were conducted by Azenta Life Sciences. Further analysis, including clustering, GSEA, and hallmark pathway analysis were performed using GSEA (v4.3.2) Mac App, iDEP 2.01, and ClusterProfiler 4.0, respectively. Briefly, Total RNA from treated and control samples (human cell lines; murine tumors) was sequenced, aligned with STAR, and quantified with DESeq2 in iDEP 2.01 ^89^. The full ranked gene list (log₂-fold change × –log₁₀ P) served as input for pre-ranked GSEA (v4.3.2) against the 50 MSigDB Hallmark sets; 1 000 permutations, FDR q < 0.25, and |NES| > 1.5 defined significance. Enrichment was independently validated with clusterProfiler 4.0 ^90^ in R, yielding concordant Hallmark pathways.

### Colony formation assay

Briefly, ZR-75-1 cells were seeded at a density of 500 cells per well in a six-well plate and allowed to grow for 24 h. The cells were then treated with drugs and incubated for 10 days at 37 °C with 5% CO_2_. After the incubation, the cell colonies were fixed in methanol, stained with crystal violet, and imaged using an AI600 Chemiluminescent Imager (Amersham).

### TUNEL assay

Detection of apoptosis were performed by the TUNEL reaction, using the TUNEL assay kit (#25879, Cell signaling technology). Cells were plated onto coverslips, and 2 days after drug treatment, they were fixed in 4% formaldehyde for a duration of 20 minutes at 24 °C. After fixation, the cells were permeabilized in 0.1% Triton X-100 for 5 minutes. The coverslips were then rinsed three times with PBS and stained with TUNEL reaction mix following the manufacturer’s instruction. The image data were analyzed under a fluorescence microscope. Experiments were evaluated in triplicate, and 10 fields of view were quantified for each sample. The coverslips were then mounted onto glass slides using ProLong™ Diamond Antifade Reagent (#P36961, ThermoFisher Scientific). Imaging was performed on a spinning disk confocal microscope (a Nikon Ti2 with a Yokogawa CSU-W1 spinning disk head) with a 20×/0.75 NA Plan Fluor Air.

### Enhanced analysis of correlation between cytokine and immune cell profiles in breast cancer samples

We first confirmed upregulation of key cytokines (CCL5, CXCL10, IFI44, IFI44L, and STAT1) involved in immune cell recruitment ^36^ due to the combination treatment. We employed the xCell algorithm (a gene signature-based method) ^37^ to analyze the association between upregulated cytokines and immune cell infiltrates in breast cancer samples.

### Multichannel fluorescence intravital microscopy (MFIM)

To visualize tumor cells in live imaging, 3.5×10^6^ MC4-L2-GFP cells were prepared for each mouse and resuspended in 100 μl of PBS. Cells were injected into the fourth mammary fat pad of 8-week-old BALB/C (Charles River Laboratories) mice. When tumors reached approximately 200 mm^3^, a combination treatment of emricasan and S64315 was administered for five consecutive days as previously described. Imaging was performed a day after the final treatment using the integrated imaging platform of MFIM (IVM-CMS3, IVIM Technology, Korea). Dendritic cells and CD8 cells were labeled by i.v. injection of FSD555 (Bioacts, #KOSC1003)-conjugated anti-CD11b antibody (BD, #553308, clone M1/70), FSD647 (Bioacts, #KOSC1315)-conjugated anti-mouse CD8a antibody (Biolegend, #100801, clone 5H10-1), respectively. Fluorescence conjugation was performed by IVI TagTM Labeling kit (IVIM Technology, Korea).

### CASP3-focused pan-cancer and immunotherapy-response analyses

First, differential expression across tumour and normal tissues was quantified with TNMplot v2.0 ^91^, which combines TCGA and GTEx RNA-seq data; log₂-TPM + 1 values were compared by Mann–Whitney test, and p < 0.05 was deemed significant. Prognostic impact was evaluated in the breast-cancer subset of KM-Plotter ^92, 93^: patients were dichotomised at the median CASP3 level, Kaplan–Meier curves for overall survival were generated, and hazard ratios with 95 % confidence intervals were calculated; a log-rank p < 0.05 indicated significance. To link CASP3 to the tumour immune milieu, we employed TIMER 2.0 ^69, 94^, which infers immune-cell infiltration from bulk RNA-seq; Spearman correlations between CASP3 expression and estimated abundances of CD8⁺ T cells, dendritic cells, and other effectors were extracted, with |ρ| > 0.3 and p < 0.05 considered meaningful. Finally, potential predictive value for checkpoint blockade was tested using ROCplotter ^95^, analysing 1,434 anti-PD-1–treated tumours from 19 datasets: CASP3 z-scores in responders (CR + PR) versus non-responders (SD + PD) were compared by Wilcoxon rank-sum test, adopting p < 0.05 as the significance threshold.

### Statistical analysis

Statistical analyses were performed using GraphPad Prism software. Data are presented as mean ± standard error (*in vivo* and *in vitro*). Appropriate statistical tests, including two-tailed unpaired t-tests, one-way ANOVA, and two-way ANOVA were used based on experimental design.

## Data availability

Data supporting the findings of this study are included within the article and its supplementary materials. Source data are available in a Source Data file.

## Acknowledgments

We would like to thank Dana-Farber Flow Cytometry core for assistance with cell sorting and Harvard Center for Biological Imaging for assistance with IHC slide scanning. We would also like to thank Dr. Richard Schwartz and Dr. Qin Yan for cell lines and Dr. Barclay Lee from DFCI Editorial Support Program for assistance with manuscript proofreading. A.S. was supported by the Burroughs-Wellcome Fund Career Award for Medical Scientists, Claudia Adams Barr Program in Innovative Cancer Research, Saverin Breast Cancer Research Grant, and JCRT Foundation grant.

---

**Extended Data Figure 1:**
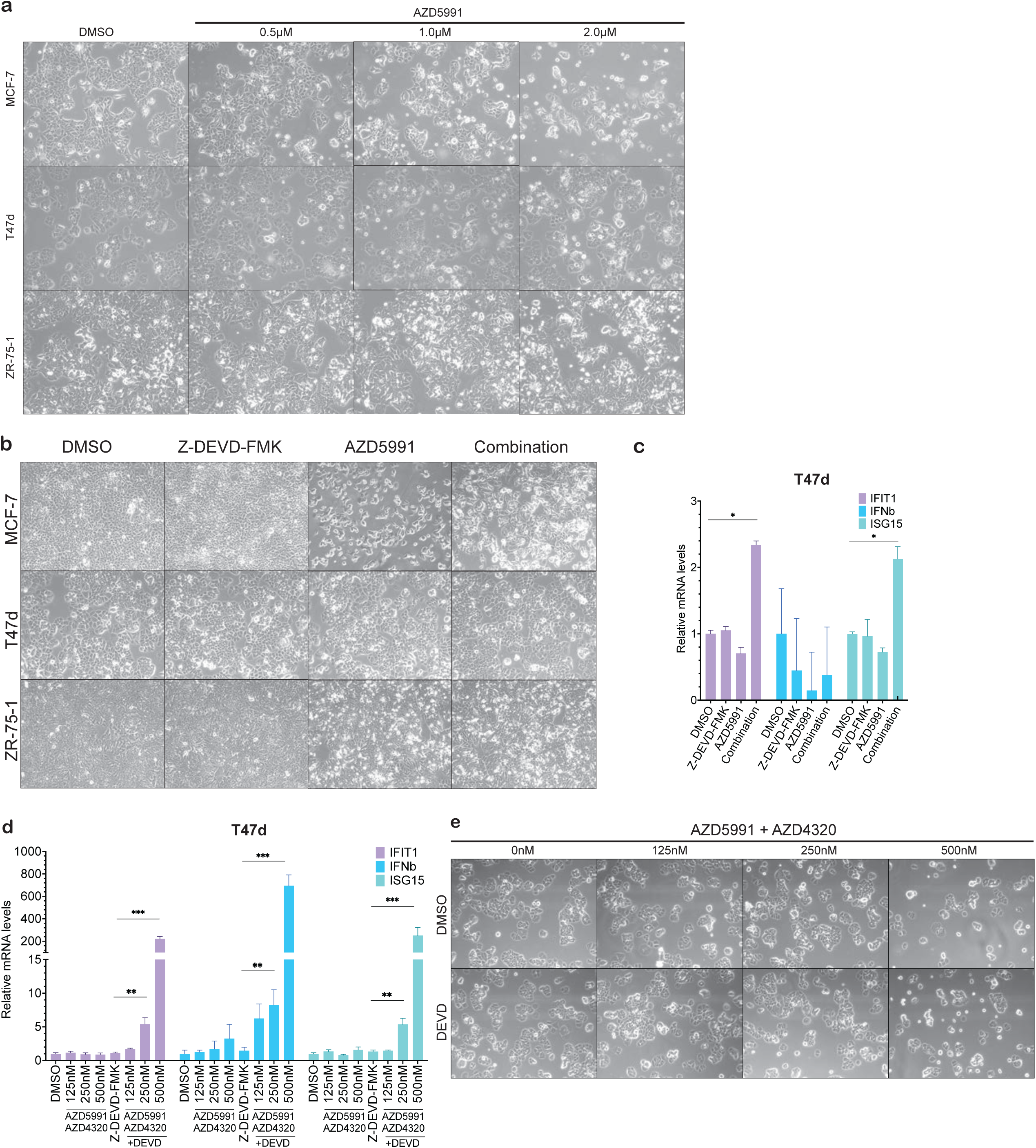
Cytotoxic and IFN response of MCL1 Inhibition alone and in combination with BCL2 Inhibition in ER+ breast cancer cell lines. **a,** Representative bright-field images (n=3 biologically independent samples) of ER+ breast cancer cell lines (MCF-7, T47d, and ZR-75-1) treated with MCL1 inhibitor AZD5991 (0.5 µM, 1.0 µM, and 2.0 µM) **b**, Representative brightfield images of MCF-7, T47d and ZR-75-1 upon the treatment with MCL1 inhibitor AZD5991 (2 µM) and/or caspase-3 inhibitor Z-DEVD-FMK (10 µM). **c,** Quantitative PCR to measure expression of IFNB1 and ISGs (IFIT1 and ISG15) upon combined treatment with MCL1 inhibitor AZD5991 (2 µM) and caspase-3 inhibitor Z-DEVD-FMK (10 µM) in T47d **d**, Quantitative PCR analysis of type I IFN genes (IFNB1, IFIT1, and ISG15) in T47d cells following treatment with MCL1 inhibitor AZD5991 (2 µM) and BCL2 inhibitor AZD4320 at varying concentrations (125nM, 250nM, and 500nM), with or withour caspase-3 inhibitor Z-DEVD-FMK (10 µM). Data represent mean ± SEM; unpaired t-test; n=3 biologically independent samples. **e,** Representative bright-field images (n=3 biologically independent samples) of T47d treated with AZD5991 and BCL2 inhibitor AZD4320 at varying concentrations (125nM, 250nM, and 500nM). Statistical significance is noted as *p < 0.05, **p < 0.01, ***p < 0.001.

**Extended Data Figure 2.**
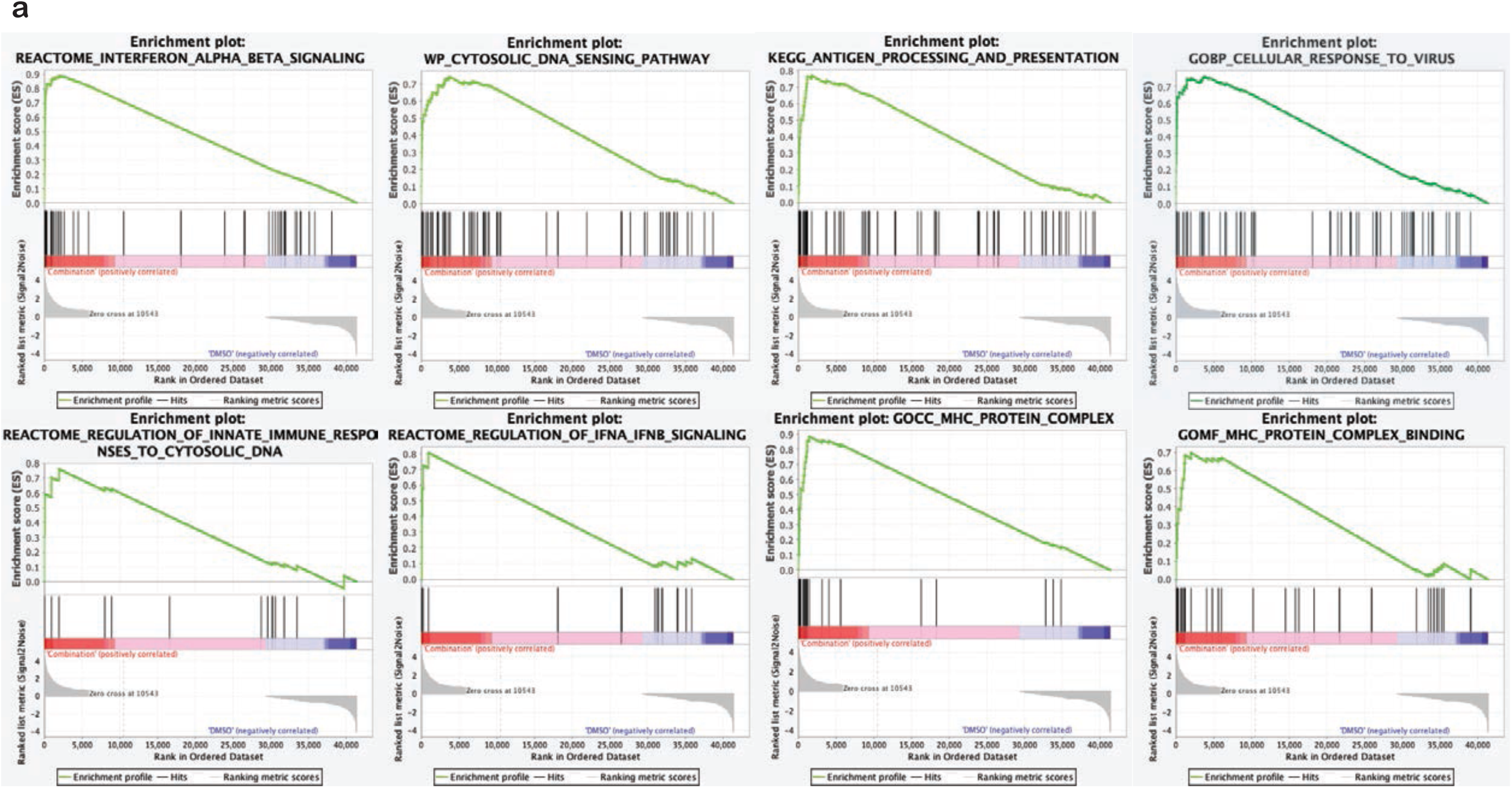
Transcriptomic alterations and pathway enhancements following combinatorial treatment in breast cancer cells. Gene Set Enrichment Analysis (GSEA, 4.3.2) plots showing significant pathway enrichments in ZR-75-1 cells treated with the combination.

**Extended Data Figure 3.**
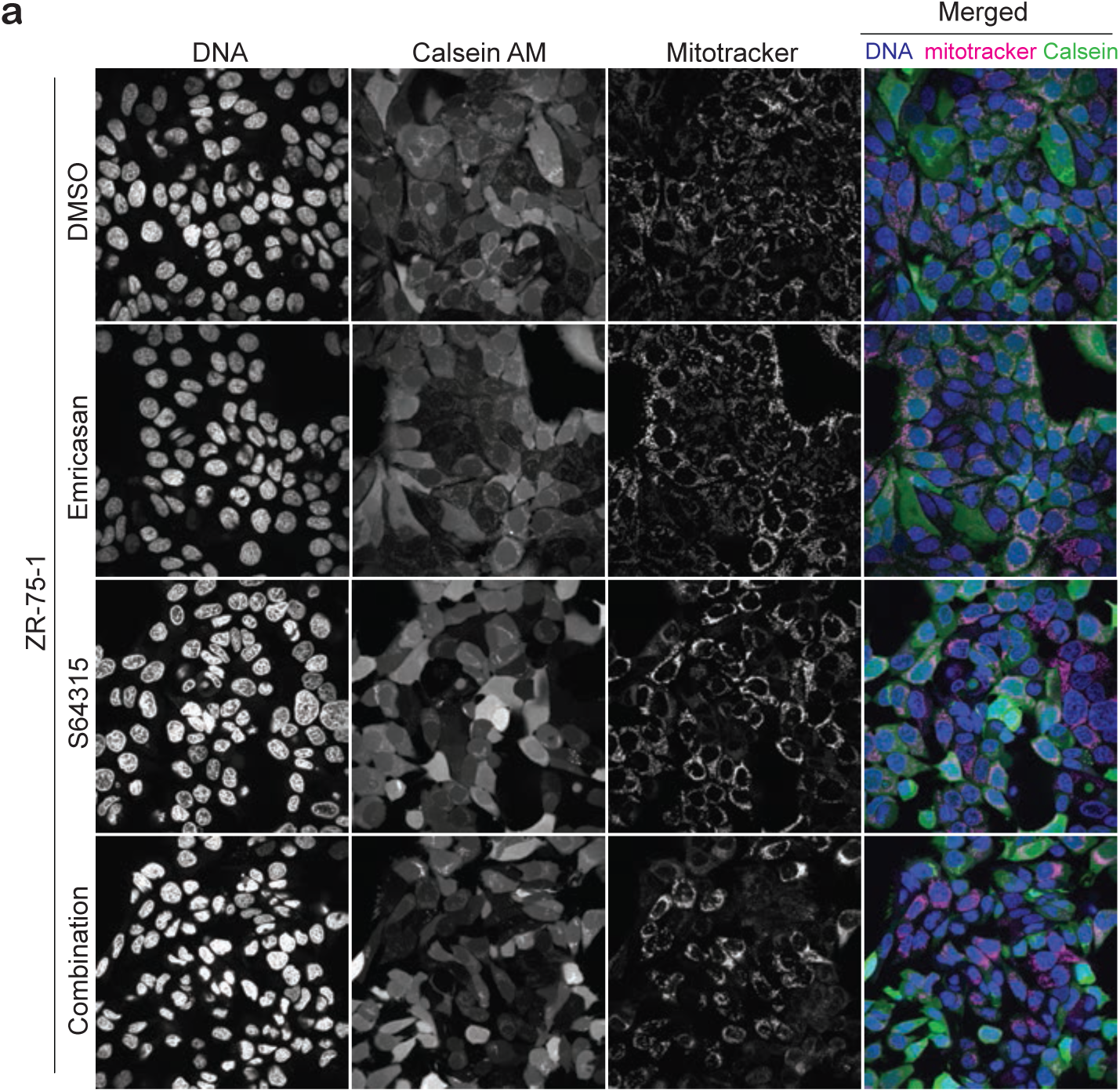
Mitochondrial permeability alteration upon the combination treatment of MCL1 and caspase inhibitors. **a,** Representative live-cell imaging analysis of ZR-75-1 cells showing DNA (blue), MitoTracker™ Red FM (magenta), and calcein AM (green). Cells were treated with a combination of MCL1 inhibitor S64315 (2 µM) and pan-caspase inhibitor Emricasan (10 µM).

**Extended Data Figure 4.**
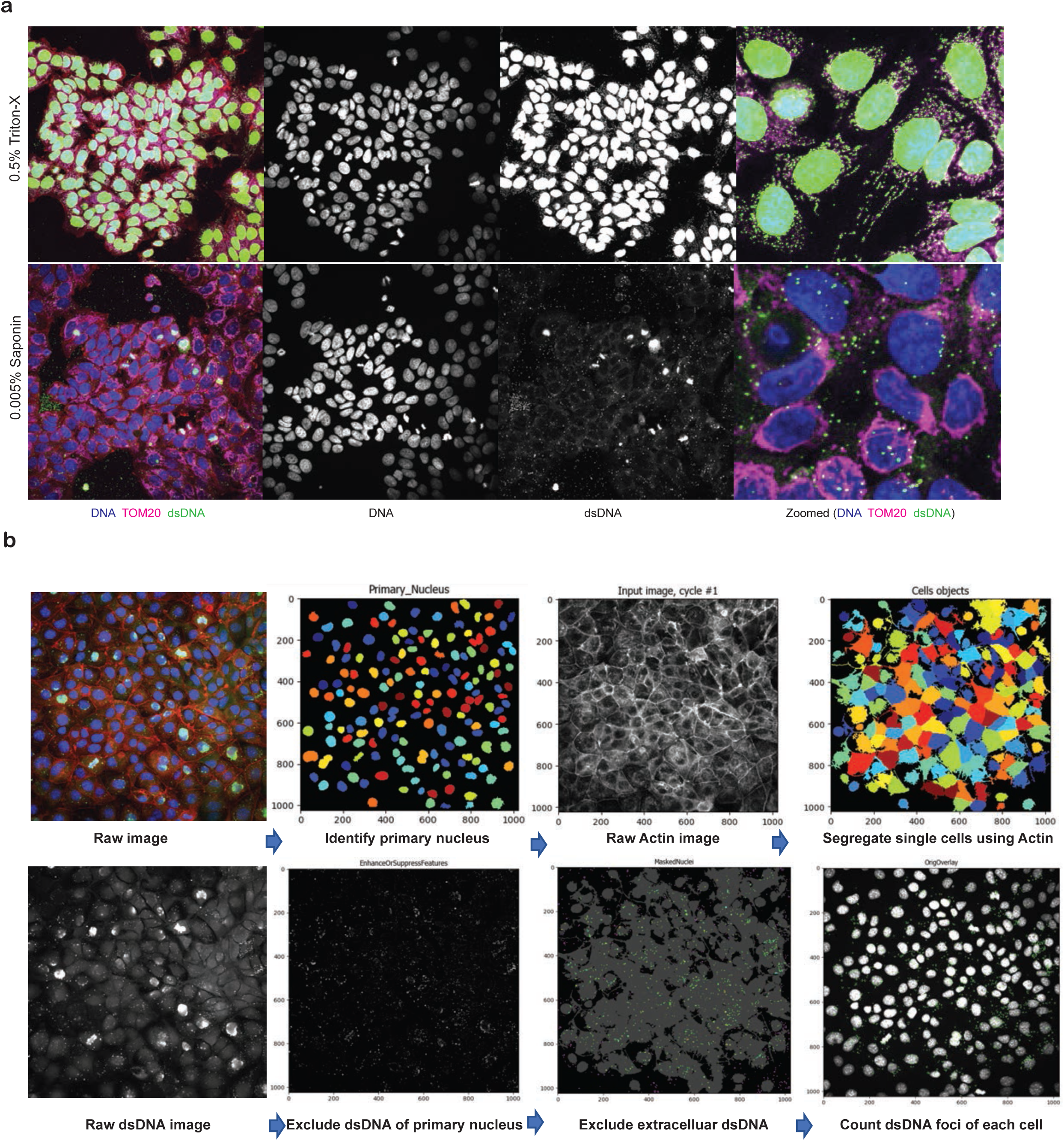
Effects of different detergents on membrane permeabilization. **a,** Representative immunofluorescence analysis (n=3 biologically independent samples) for dsDNA (green), the mitochondrial marker TOM20 (magenta), and cell nuclei (DNA, blue) in cells pre-treated with either 0.5% Triton-X or 0.005% saponin to permeabilize different cellular membrane. **b**, Representation of the quantification process using CellProfiler (Version 4.2.6).

**Extended Data Figure 5.**
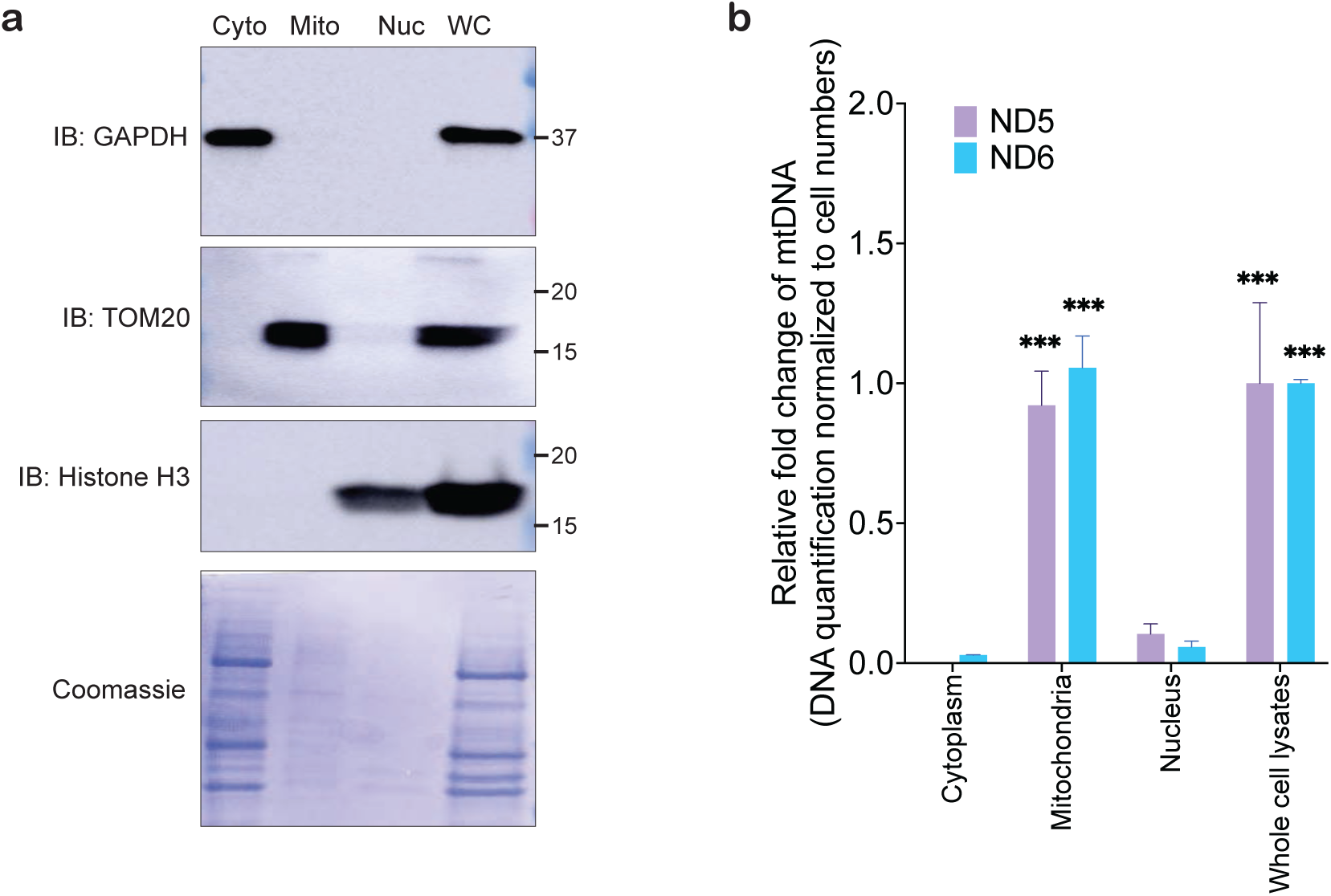
Confirmation of mitochondrial DNA compartmentalization a, Western blot analysis and Coomassie staining in fractionated cell lysates, with GAPDH as a cytoplasmic marker, TOM20 as a mitochondrial marker, and Histone H3 as a nuclear marker, across cytoplasmic (Cyto), mitochondrial (Mito), nuclear (Nuc), and whole-cell (WC) fractions. (n=3 biologically independent samples) **b,** Quantitative PCR analysis of mitochondrial DNA (mtDNA) content for genes ND5 and ND6 in different cellular compartments of MCF7 cells treated with the MCL1 inhibitor alone or in combination with the caspase inhibitor. Data represent mean +/- SEM; unpaired t-test; n=3 biologically independent samples. Statistical significance is noted as *p < 0.05, **p < 0.01, ***p < 0.001.

**Extended Data Figure 6.**
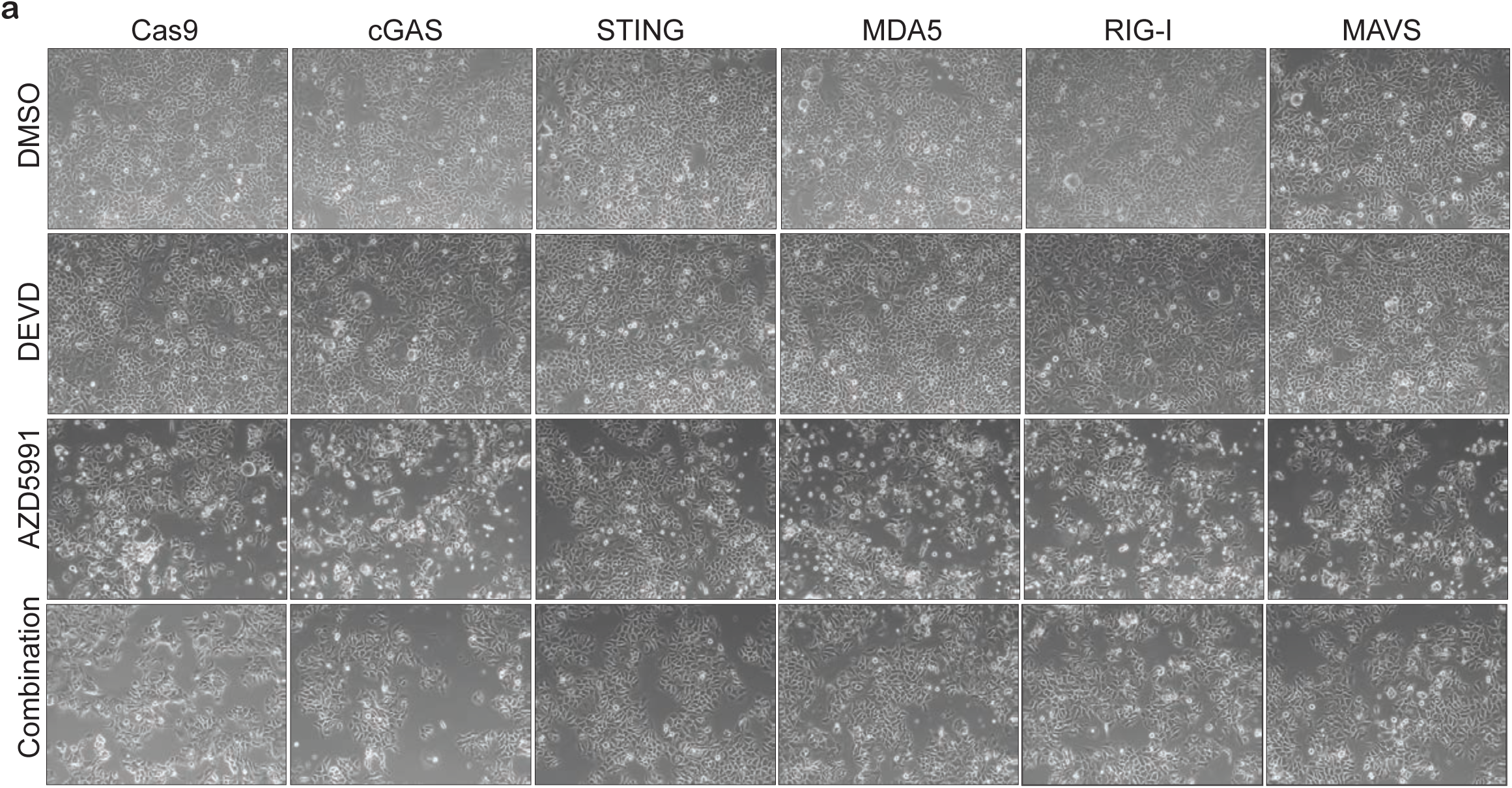
The cytotoxicity assessment in MCF7 CRISPR/Cas9 knockout cell lines. **a,** Representative bright field images (n=3 biologically independent samples) of MCF7 CRISPR/Cas9 knockout cell lines for cGAS, STING, MDA5, RIG-I, and MAVS, treated with DMSO, the caspase inhibitor Z-DEVD-FMK (2 µM), the MCL1 inhibitor AZD5991 (10 µM), and their combination.

**Extended Data Figure 7.**
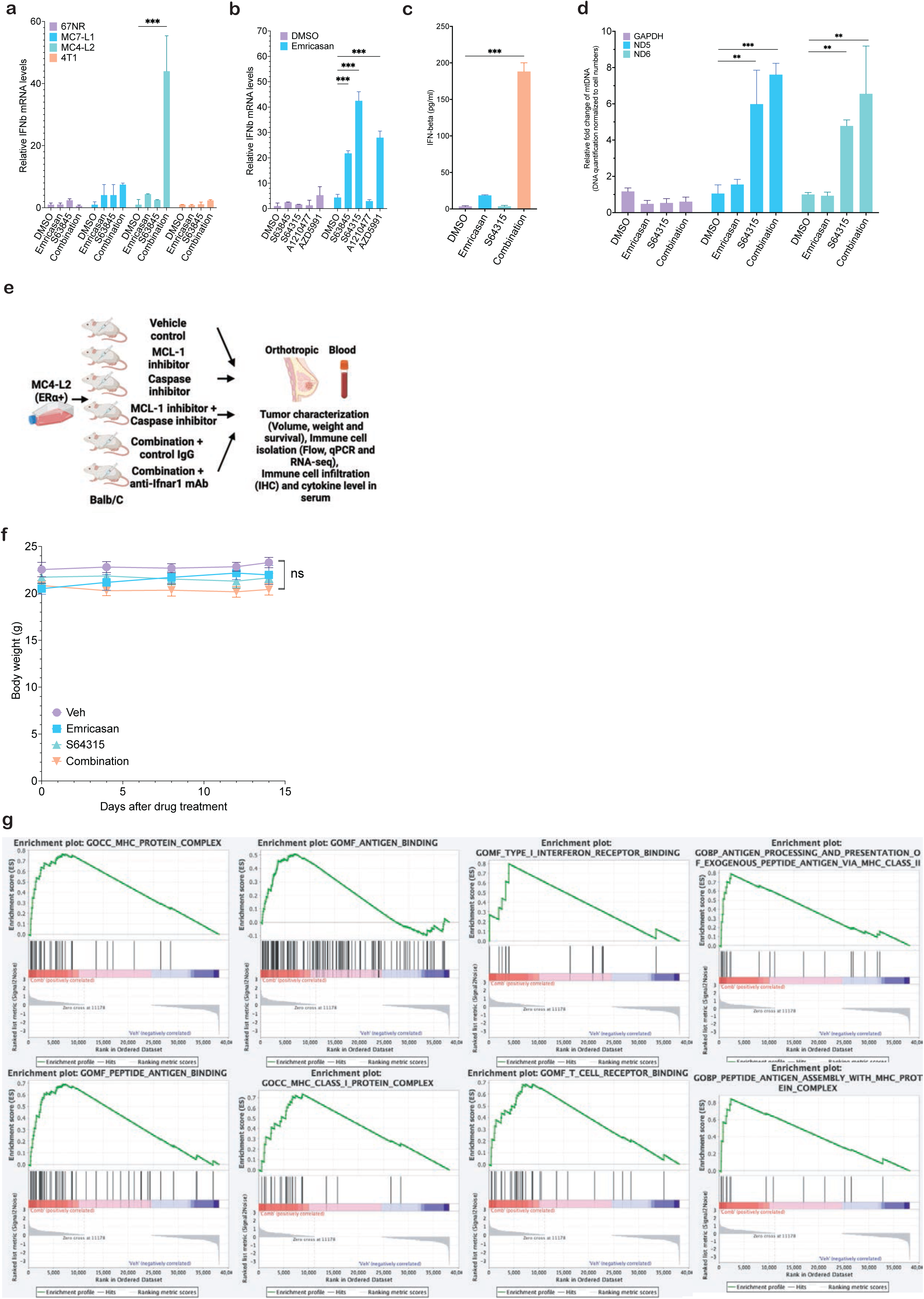
Pre-evaluation of IFN response in murine breast cancer cell lines, toxicity analysis in murine tumor models, and transcriptomic alterations following combinatorial treatment. **a,** Quantitative PCR analysis of IFNB1 mRNA levels in various murine breast cancer cell lines, including 67NR, 4T1, MC7-L1, and MC4-L2, treated with MCL1 inhibitor (2 µM S64315), caspase inhibitor (10 µM emricasan), and their combination. Data represent mean ± SEM; unpaired t-test; n=3 biologically independent samples. **b,** Quantitative PCR analysis of IFNB1 in MC4-L2 cells after treatment with different MCL1 inhibitors (2 µM S63845, 2 µM S64315, 2 µM A1210477, 2 µM AZD5991) in combination with emricasan (10 µM). Data represent mean ± SEM; unpaired t-test; n=3 biologically independent samples. **c,** ELISA analysis of IFN-β levels in media of MC4-L2 after treatment with MCL1 inhibitor (2 µM S64315), caspase inhibitor (10 µM emricasan), and their combination. Data represent mean ± SEM; unpaired t-test; n=3 biologically independent samples. **d,** Quantitative PCR analysis of mitochondrial DNA (mtDNA) genes ND5 and ND6 in cytoplasmic fractions of MC4-L2 upon the treatment with MCL1 inhibitor S64315 (2 µM) and/or caspase inhibitor emricasan (10 µM). Data represent mean +/- SEM; unpaired t-test; n=3 biologically independent samples. **e,** Treatment group and analysis plan of *in vivo* study. **f,** Murine breast cancer cell line (MC4-L2) was injected into the mammary fat pad of wildtype mice (BALB/c) treated with S64315 and/or emricasan. **g,** Gene Set Enrichment Analysis (GSEA, 4.3.2) plots showing significant pathway enrichments in mice treated with the combination. Analysis of body weight changes of MC4-L2 tumors in mice (Veh, n=5; emricasan, n=4; S64315, n=6; Combination, n=8). Data represent mean +/- SEM; two-way ANOVA. Data represent mean +/- SEM; two-way ANOVA. Statistical significance is noted as *p < 0.05, **p < 0.01, ***p < 0.001 and ns, statistically not significant.

**Extended Data Figure 8.**
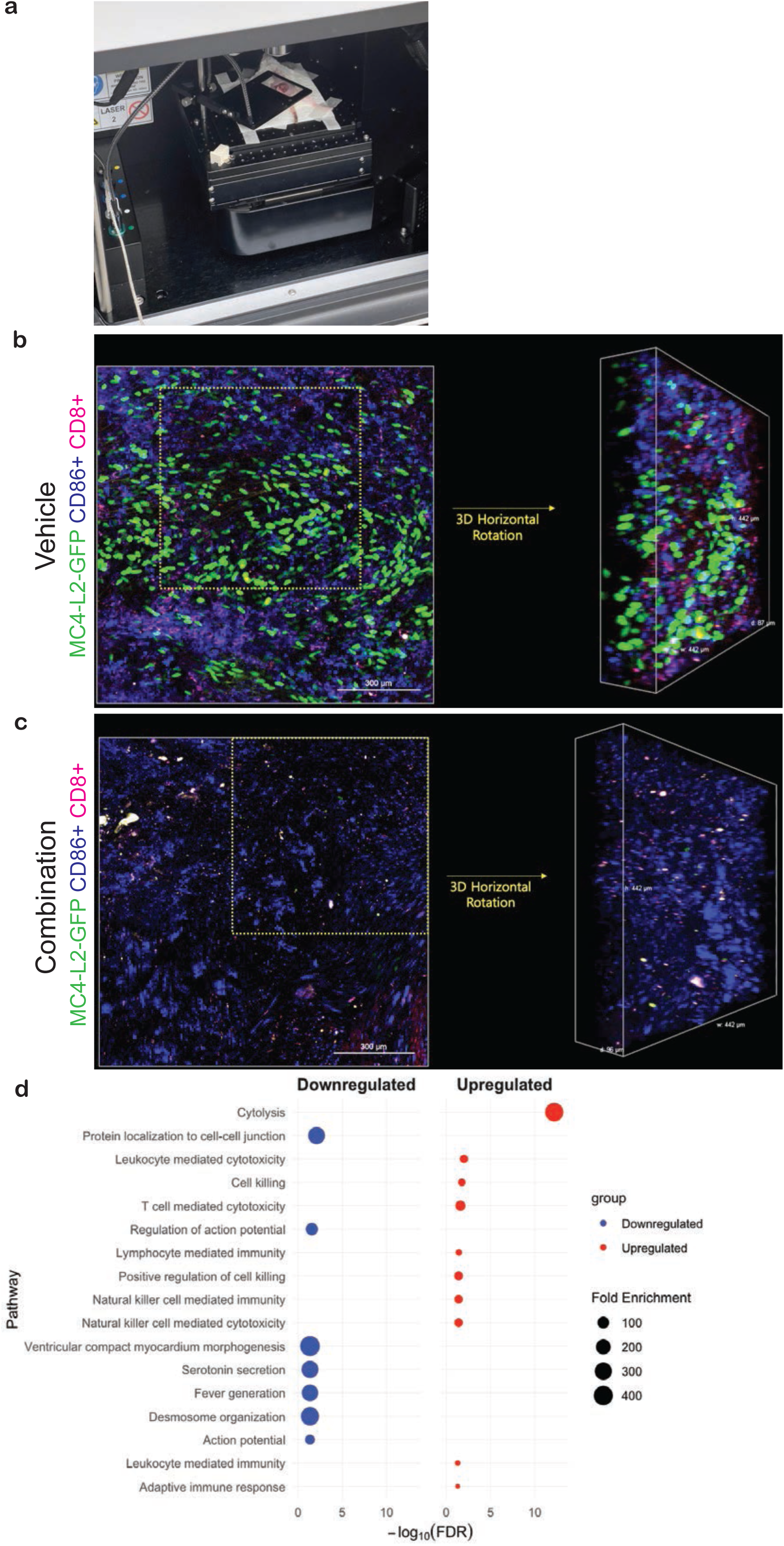
*In-vivo* imaging and 3D visualization of MC4-L2-GFP tumors treated with vehicle or combination therapy. **a**, Imaging setup used for in-vivo imaging of MC4-L2-GFP tumors. **b and c**, Representative images and 3D reconstructions of MC4-L2-GFP tumors in BALB/C mice treated with either vehicle (top) or a combination of S64315 and emricasan (bottom). Tumors were imaged immediately after 5 consecutive days of treatment. The left column shows 2D images, with green fluorescence indicating MC4-L2-GFP cells, magenta representing CD86+ dendritic cells, and red indicating CD8+ T cells. The yellow boxes highlight regions of interest, which are further analyzed in the 3D horizontal rotation views (right column). Scale bars: 300 µm (2D images) and 442 µm (3D images). **d**, Top 17 GO gene sets by iDEP DEGs (Vehicle vs combination treatment, n=4 mice for each treatment condition).

**Extended Data Figure 9.**
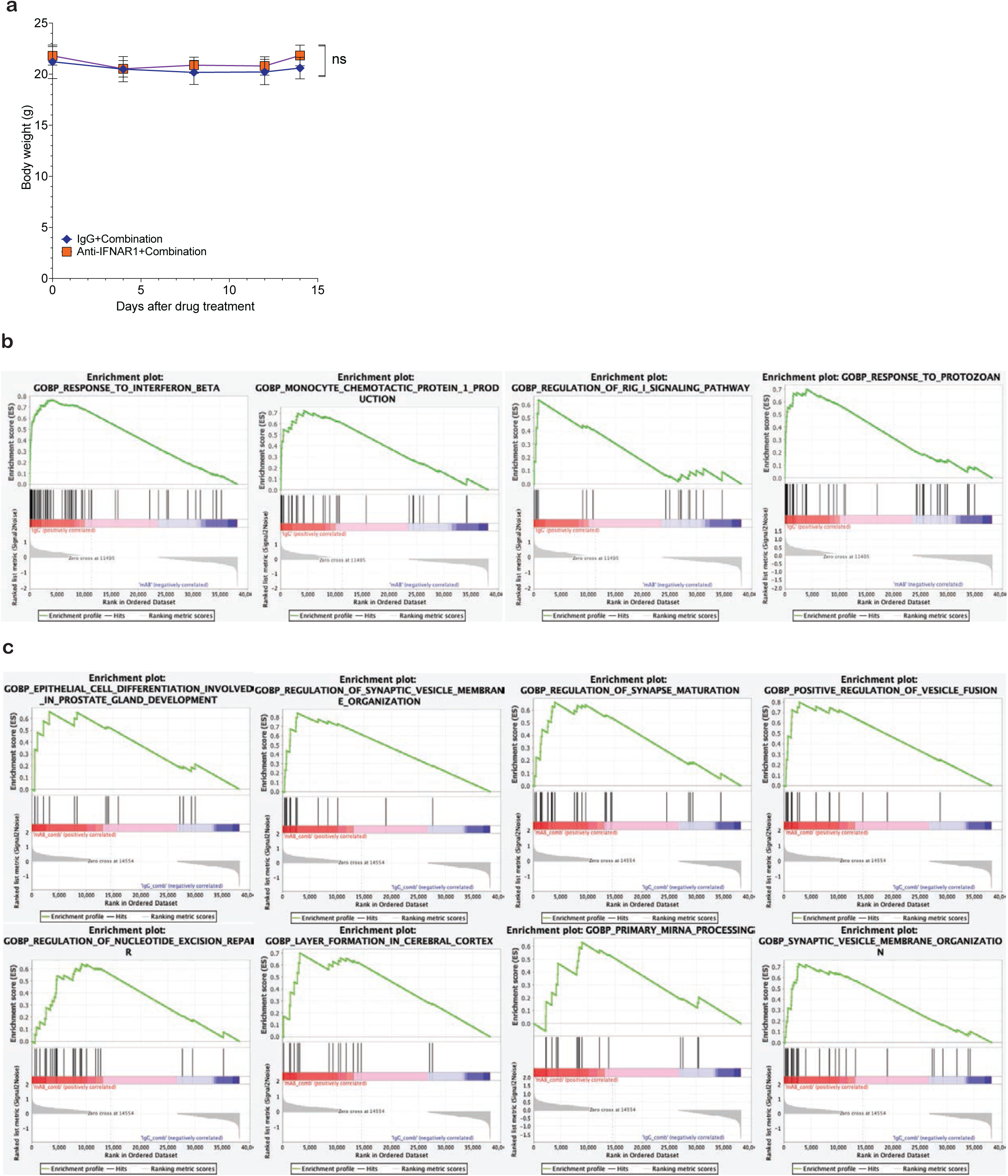
IFNAR1-dependent toxicity and transcriptomic responses induced by MCL1 and caspase inhibitor combination therapy in murine breast cancer models treated with control IgG or anti-IFNAR1 antibody. **a,** Murine breast cancer cell line (MC4-L2) was injected into the mammary fat pad of wildtype mice (BALB/c) treated with S64315 and emricasan. Prior to these treatments, mice were pre-treated with either IgG or anti-IFNAR1 monoclonal antibody. Analysis of body weight changes of MC4-L2 tumors in mice (IgG+combination, n=7; anti-IFNAR1+combination, n=8). Data represent mean +/- SEM; two-way ANOVA. Statistical significance is noted as *p < 0.05, **p < 0.01, ***p < 0.001 and ns, statistically not significant. **b,** Gene Set Enrichment Analysis (GSEA, 4.3.2) plots showing significant pathway enrichments in mice treated with Control IgG and the dual combination. **c,** Gene Set Enrichment Analysis (GSEA, 4.3.2) plots showing significant pathway enrichments in mice treated with anti-IFNAR1 monoclonal antibody and the dual combination.

**Extended Data Figure 10.**
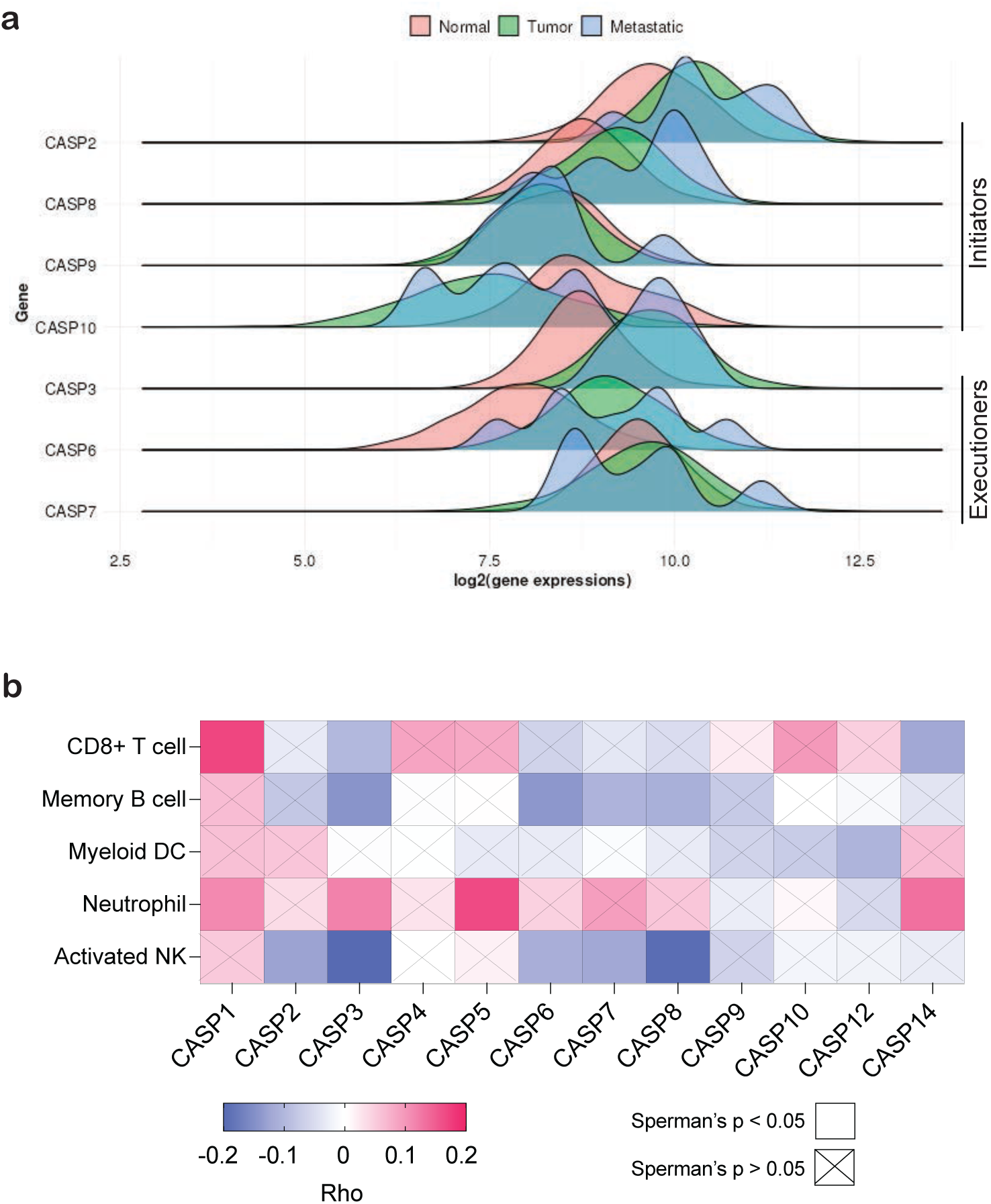
Correlation of CASP3 expression with survival outcomes, immune infiltration and impaired immunotherapy response in breast cancer. **a**, Expression levels of caspases (CASP2, CASP3, CASP6, CASP7, CASP8, CASP9, and CASP10) plotted with https://tnmplot.com/ webtool ^91^. **b**. Kaplan-Meier plots based upon CASP3 expression levels plotted with the kmplot.com webtool ^96^. **c**, Heatmap generated by timer.cistrome.org webtool showing Spearman correlation of 12 caspase expression with infiltration scores for CD8+ T cels, memory B cell, myeloid DC, neutrophils and activated NK cells. Color code shows level of correlation from +1 (positive correlation) to 1 (negative correlation) by Rho and significance indicated by the p value ^69, 94, 97^. **d**, Infiltration level of memory B cells and activated NK cells in *in vivo* RNAseq data analyzed by timer.cistrome.org webtool. Data represent mean +/- SEM; unpaired t-test, n=4 biologically independent samples. **e**, Box plot showing distribution of CASP3 signature score in 1434 tumor tissue samples from 19 datasets with esophageal, gastric, head and neck, lung, and urothelial cancers, plus melanoma, separated by responders and non-responders to anti-PD-1 (nivolumab, pembrolizumab) agents using rocplot.com webtool ^98^.

**Supplementary movie 1.** Representative live-cell imaging of ZR-75-1 cells expressing BAF-GFP and H2b-RFP treated with DMSO. Images were captured every 15 minutes over a 48-hour period. Color legend: Green-BAF; Red-H2b; Magenta-Mitotracker.

**Supplementary movie 2.** Representative live-cell imaging of ZR-75-1 cells expressing BAF-GFP and H2b-RFP treated with a combination of MCL1 inhibitor S64315 (2 µM) and pan-caspase inhibitor Emricasan (10 µM). Images were captured every 15 minutes over a 48-hour period. Color legend: Green-BAF; Red-H2b; Magenta-Mitotracker.

